# The AMPK-related kinase NUAK1 controls cortical axons branching though a local modulation of mitochondrial metabolic functions

**DOI:** 10.1101/2020.05.18.102582

**Authors:** Marine Lanfranchi, Géraldine Meyer-Dilhet, Raphael Dos Reis, Audrey Garcia, Camille Blondet, Luc Javin, Alizée Amar, Julien Courchet

## Abstract

The precise regulation of the cellular mechanisms underlying axonal morphogenesis is essential to the formation of functional neuronal networks. We previously identified the autism-candidate kinase NUAK1 as a central regulator of axon branching in mouse cortical neurons through the control of mitochondria trafficking. How does local mitochondrial position or function regulate axon branching during development? Here, we characterized the metabolic regulation in the developing axon and report a marked metabolic decorrelation between axon elongation and collateral branching. We next solved the cascade of event leading to presynaptic clustering and mitochondria recruitment during spontaneous branch formation. Interestingly and contrary to peripheral neurons, mitochondria are recruited after but not prior to branch formation in cortical neurons. Using flux metabolomics and fluorescent biosensors, we observed that NUAK1 deficiency significantly impairs mitochondrial metabolism and axonal ATP concentration. Upregulation of mitochondrial function is sufficient to rescue axonal branching in NUAK1 null neurons *in vitro* and *in vivo*. Altogether, our results indicate that NUAK1 exerts a dual function during axon branching through its ability to control mitochondria distribution and activity, and suggest that a mitochondrial-dependent remodeling of local metabolic homeostasis plays a critical role during axon morphogenesis.

## INTRODUCTION

In mammals, cortical circuits are organized into complex networks of neurons interconnected through long-range axonal projections. The establishment of circuit connectivity occurs during a protracted period of embryonic and postnatal development through a series of tightly regulated cellular processes whose disruption can lead to socially-devastating neurodevelopmental disorders such as autism spectrum disorders (ASD), schizophrenia, epilepsy or mental retardation (del Pino et al., 2018; Piven et al., 2017). Axon morphogenesis is a multistep process culminating with the formation of axonal branches to form a network that is later refined through a process of selection of functional contacts and elimination by pruning (Kalil and Dent, 2014; Lewis et al., 2013). The formation and refinement of terminal axonal branches relies on a balance between synaptic activity and target-derived extracellular neurotrophins (Mizuno et al., 2007; C.-L. Wang et al., 2007). These signals converge on the activation of intracellular signaling pathways mediating a cascade of cellular events such as cytoskeleton remodeling, addition of membrane or local protein translation (Brosig et al., 2019; Dent et al., 1999; Dent and Kalil, 2001; Hu et al., 2012; Ponomareva et al., 2014; A. Schwarz et al., 1995; Spillane et al., 2012; 2013; Wong et al., 2017) that are highly taxing energetically and are thought to induce a local increase in the metabolic turnover. Yet the spatial diffusion of metabolic molecules such as ATP is limited, implying that cellular energy production must match the local demand. Understanding how these local mechanisms are set up and how they contribute to aspects of neuronal development such as the shaping of axonal complexity is therefore a key question in cellular neuroscience.

The local metabolic activity has been best studied at synapses (Lee et al., 2018; Rangaraju et al., 2019; Rossi and Pekkurnaz, 2019), which account for the majority of ATP consumption in neurons to ensure ionic homeostasis and synaptic vesicle recycling (Attwell and Laughlin, 2001; Harris et al., 2012; Rangaraju et al., 2014; Vos et al., 2010). In agreement with this local metabolic need, mitochondria are actively transported to and captured at presynapses (J. Courchet et al., 2013; Devine and Kittler, 2018; Hollenbeck and Saxton, 2005; Misgeld and T. L. Schwarz, 2017). Mitochondrial function has been proposed to support synaptic release through a crosstalk between ATP-producing oxidative phosphorylations (Ashrafi et al., 2020; Rangaraju et al., 2014) and calcium buffering (Hirabayashi et al., 2017; Kwon et al., 2016; Vaccaro et al., 2017; Villegas et al., 2014), converging on the fine-tuned regulation of synaptic vesicle release. In contrast, the regulation of local metabolism in immature neurons and during axon morphogenesis is far less well-understood. Recent studies uncovered that mitochondria function is critical for neuronal development and cortical circuits formation (Fernandez et al., 2019; Klein Gunnewiek et al., 2020; Oruganty-Das et al., 2012).

Deregulation of mitochondria positioning, morphology or function disrupts the formation of axonal projections *in vivo* (J. Courchet et al., 2013; Fernandez et al., 2019; Lewis et al., 2018). In developing retinal and sensory axons, mitochondria function has been linked to the local protein synthesis required for cytoskeleton remodeling and the conversion of filopodia into a collateral axonal branch (Cioni et al., 2019; Spillane et al., 2013; Wong et al., 2017). However it is unclear if all neuronal subtypes or if all modalities of axonal branching rely on the same mechanisms for branch formation, growth or consolidation. Hence there is a need to better understand the cellular mechanisms underlying the role of mitochondrial function in the formation and refinement of axon morphogenesis and thereby the establishment of circuit connectivity.

We previously identified that a signaling pathway formed by the kinases LKB1 (STK11) and NUAK1 (ARK5) control cortical axon branching through the regulation of mitochondria capture at immature presynaptic sites (J. Courchet et al., 2013). NUAK1 belongs to a family of 14 closely related kinases related to the metabolic regulator AMPK (PRKAA) defining the so-called AMPK-related kinases (AMPK-RK) family (Lizcano et al., 2004). NUAK1 expression is strongly enriched in the developing mouse cortex (Hirano et al., 2006; Ohmura et al., 2012). Rare *de novo* mutations of NUAK1 have been associated to several neurodevelopmental disorders including Autism Spectrum Disorders (ASD) (Iossifov et al., 2014; 2012), Attention Deficit / Hyperactivity Disorders (ADHD) (Alemany et al., 2015), cognitive impairment (Johnson et al., 2016) or hydrocephaly (Vojinovic et al., 2018). In mice, NUAK1 heterozygosity affects cortical development, leading to an array of anatomical and behavioral deficits (V. Courchet et al., 2018). So far, the cellular functions of NUAK1 remain incompletely known. It has been linked to the control of the actomyosin cytoskeleton through the phosphorylation of the Protein Phosphatase 1 (PP1) regulatory subunit MYPT1 (Zagórska et al., 2010). Furthermore NUAK1 directly phosphorylates the microtubule-associated protein TAU (Lasagna-Reeves et al., 2016) and accordingly NUAK1 inhibition can alleviate some of the cellular and behavioral effects in a mouse model of tauopathy. In parallel NUAK1 function has been tied to the regulation of mitochondrial metabolism in Myc-overexpressing cancer cells, who rely on NUAK1 expression to maintain metabolic homeostasis and for survival (Liu et al., 2012).

Our previous observation that axonal mitochondria capture at nascent presynaptic boutons is essential for terminal axon branching (J. Courchet et al., 2013) raised a central and unresolved question pertaining to the function of presynaptic mitochondria in axon branching. In this study, we explored the hypothesis that presynaptic mitochondria have a unique property with regard to their ability to promote branch formation or stabilization in cortical pyramidal neurons (PNs). We observed that axon elongation and collateral branching rely on distinct metabolic pathways. We further characterized that mitochondria accumulate specifically at presynaptic sites located at axonal branch points, and that unlike presynaptic boutons that are formed before axonal branches, mitochondria recruitment occurs after the onset of branch formation. Through mitochondria photoinactivation, we confirmed that in cortical PNs, mitochondria are required for branch stabilization rather than branch initiation. We subsequently characterized mitochondrial function in NUAK1 deficient neurons and report that NUAK1 controls not only mitochondrial trafficking, but also mitochondrial metabolic activity. Finally using pharmacological and genetic approaches to boost mitochondrial fitness, we observed that a metabolic upregulation is sufficient to rescue axonal branching in NUAK1 deficient neurons *in vitro* and *in vivo*. Overall, our results indicate that the kinase NUAK1 exerts a dual effect on mitochondria trafficking and activity in cortical PNs and suggest a two-hit model by which branch stabilization relies not only on the proper localization of mitochondria, but also on their local activity.

## RESULTS

### Cortical axon elongation and collateral branching rely on distinct metabolic pathways

We first tested how axon morphogenesis rely on glucose metabolism (**Fig. 1A**). Cortical and hippocampal neuronal cultures are classically grown in a glucose-rich medium. To determine to what extent axonal elongation and collateral branching rely on glucose as a source of carbon, we cultured cortical neurons in neurobasal medium with no glucose, or with low (2.5mM), medium (5mM) and high (25mM) concentrations of glucose. We observed a marked dose-response effect of medium glucose concentration on axon development (**Fig. 1B**). Quantifications revealed that axonal length increases as a function of glucose concentration (**Fig. 1C**). In contrast, the number of collateral branches was dependent upon the presence, but not quantity of glucose (**Fig. 1D**). Interestingly a low dose of pyruvate (1mM) was also sufficient to support axonal branching, suggesting that collateral branch formation or stabilization is largely dependent upon mitochondrial metabolism.

**Figure 1:**
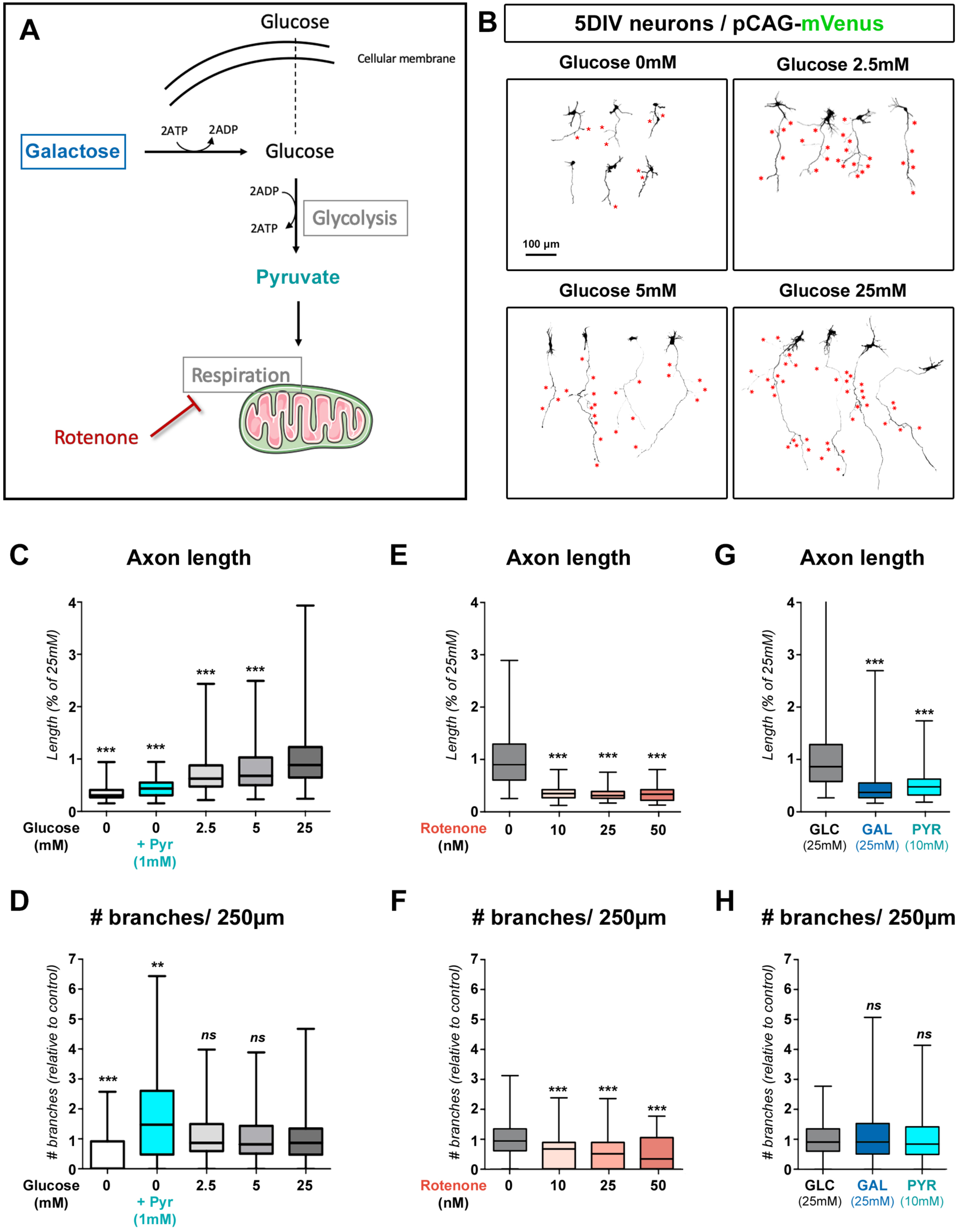
differential requirement of glycolysis and oxidative metabolism for axon growth and branching. (A) Main pathways of glucose metabolism. Galactose was used to negate the metabolic effect of glycolysis. Pyruvate was used to support mitochondrial metabolism in a glucose-free medium. Low doses of rotenone were used to lower mitochondrial Complex I activity. (B) Representative images of mVenus expressing cortical neurons (5DIV) cultured in Neurobasal medium containing the indicated doses of glucose. Red stars point to collateral branches of the axon. (C-H) Quantification of axon length and collateral branches of 5DIV neurons in the indicated conditions: (C-D) medium containing increasing concentrations of glucose. (E-F) 25mM glucose-containing medium with increasing doses of rotenone. (G-H) medium containing either glucose (25mM), or galactose (24mM) + glucose (1mM), or pyruvate (10mM) without glucose. Box-plot: 75^th^ percentile, median and 25^th^ percentile. Statistical tests: Kruskal-Wallis test with Dunn’s post-test (each condition compared to untreated 25mM condition). (C-D) N_(0mM)_=48, N_(0mM+Pyr)_=75, N_(2.5mM)_=244, N_(5mM)_=191, N_(25mM)_=238. (E-F) N_(0nM)_=120, N_(10nM)_=102, N_(20nM)_=82, N_(50nM)_=44. (G-H) N_(Glucose)_=246, N_(Galactose)_=248, N_(Pyruvate)_=130

To test this hypothesis further, we next used strategies to decouple the two main pathways of glucose catabolism, i.e. glycolysis and mitochondrial respiration (**Fig. 1A**). We used low doses of rotenone, an inhibitor of OXPHOS complex I, to downregulate mitochondrial oxidative phosphorylation capacity without affecting glycolysis. This led to a decrease of both axon length and collateral branching (**Fig. 1E-F** and **Fig. S1A**). Conversely, we bypassed glycolysis either by replacing glucose by galactose (a 6-carbon sugar catabolized by the glycolytic machinery but negating the net ATP production of glycolysis), or by adding pyruvate to a no-glucose medium. In both case, axon branching was largely equivalent to control conditions (25mM glucose) despite a strong adverse effect on axon length (**Fig. 1G-H** and **Fig. S1B-C**). Interestingly glucose concentration also affected the number of dendritic branches formed by neurons. Dendrite growth and branching was reduced, albeit slightly, in conditions where glycolysis was bypassed, whereas reduction of dendritic growth and branching was significantly more pronounced upon mitochondria inhibition by rotenone (**Fig. S1D-F**). Taken together, our results show distinct metabolic requirement for axon elongation and collateral branching and suggest that mitochondrial oxidative phosphorylation is necessary and sufficient for axon branch formation and/or consolidation.

### Mitochondria cluster at axonal branchpoint of cortical pyramidal neurons

The importance of oxidative metabolism for axonal branching prompted us to define the position of axonal mitochondria relative to collateral branches in cortical pyramidal neurons (PNs). Indeed previous studies by us and others linked branching with the trafficking and distribution of axonal mitochondria (J. Courchet et al., 2013; Sainath et al., 2017; Spillane et al., 2013). We performed *Ex Vivo* Cortical Electroporations (EVCE) of plasmids encoding the fluorescent protein BFP together with mitochondria-targeted DsRed and the synaptic vesicle marker vGlut1-GFP as a marker of immature presynaptic boutons in cortical layer 2/3 PNs mitochondria. (**Fig. 2A-B**). After 5 days of culture *in vitro* (DIV), cortical PNs were fixed and imaged to analyze the distribution of mitochondria along the axon. Mitochondria were not distributed evenly along the axon, but rather we could observe zones of high mitochondrial density and zones of low mitochondrial density (**Fig. 2A-C**). The observed distance from a given mitochondrion to its nearest neighbor was significantly lower than the expected distance if mitochondria were evenly distributed along the axon (**Fig. 2D**), indicating that mitochondria form clusters along developing axons of cortical PNs. We observed that branchpoints are significantly closer to a mitochondrion than a random position in the axon (**Fig. 2E**) and that mitochondria density was higher at branch origin than in any random location along the axon shaft (**Fig. 2F**). Accordingly, branches were found in regions of high mitochondria density more frequently than in regions of low mitochondria density (**Fig. 2G**).

**Figure 2:**
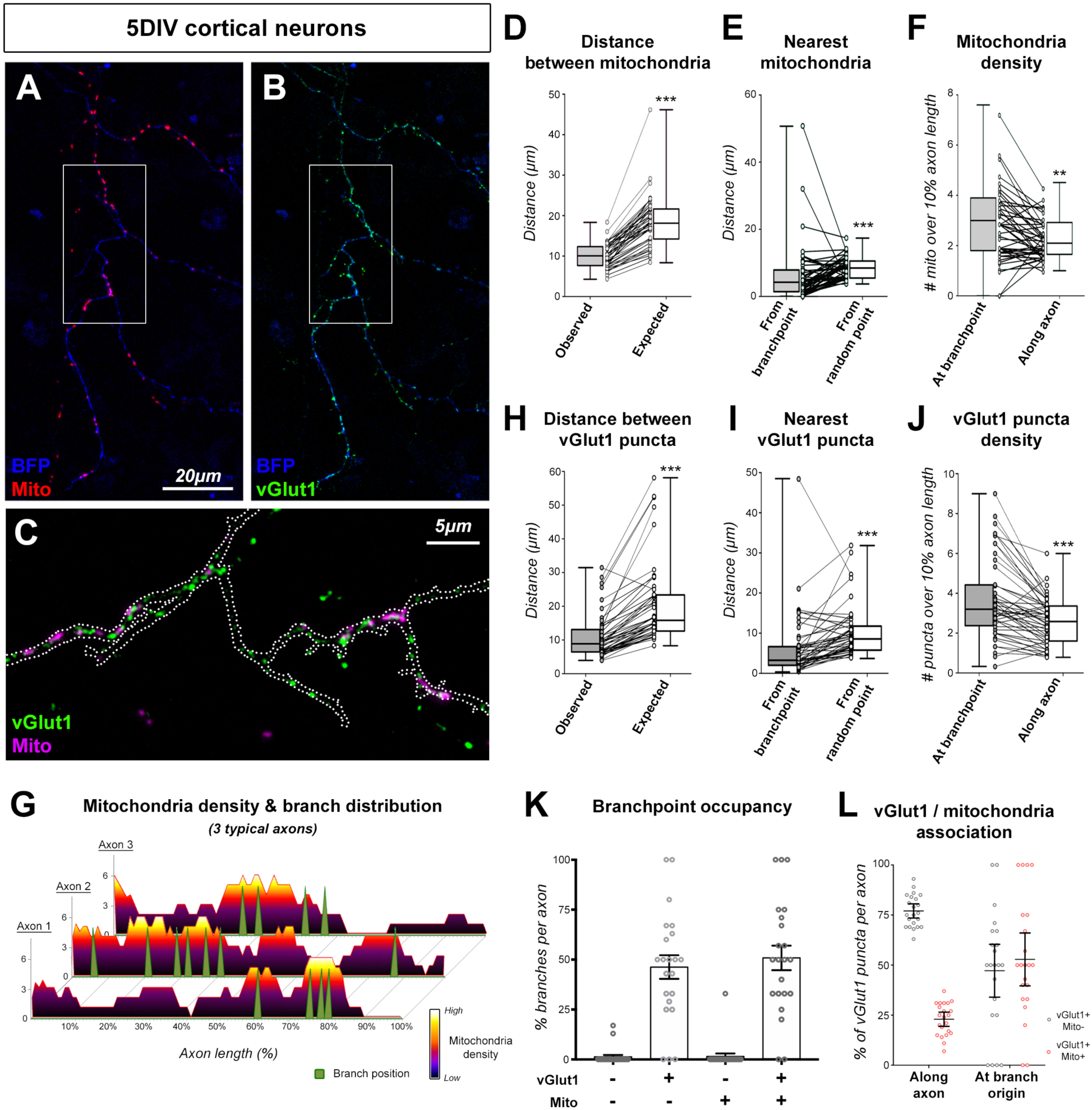
presynaptic sites and mitochondria clustering at axonal branchpoints. (A-B) Representative images of a 5DIV cortical neuron following expression of the mitochondrial marker mito-DsRed (A) or the presynaptic marker vGlut1-GFP (B). BFP was used to visualize neuronal morphology. Magnification of a segment of axon (C) shows vGlut1-GFP and mito-DsRed localization respective to branch origin (white dotted line). (D-I) Quantification of mitochondria (D-F) and presynaptic puncta (G-I) distribution in the axon. The following quantifications were performed: (D;G) observed minimal distance between objects compared to expected distance if objects were evenly distributed. (E;H) Observed minimal distance from branchpoint compared to average distance from a random point along the axon. (F;I) Object density at branchpoint compared to average density along the axon. Box-plot: 75^th^ percentile, median and 25^th^ percentile. Individual dots represent average (per axon). Statistical tests: Wilcoxon matched-pairs ranked test. N_(mito)_=46, N_(vGlut1)_=48 out of 3 independent neuronal cultures. (J) Representation of mitochondria density along the axon (normalized to axonal length from origin (0%) to extremity (100%). Position of collateral branches (green arrowhead) correlates with regions of high mitochondrial density. (K) Quantification of branchpoint occupancy by either vGlut1+ puncta, mitochondria, or vGlut1+ puncta and mitochondria. Each point represents the value (%) for a given axon. Bars: average ± SEM. N=22 axons out of 2 independent experiments. (L) Proportion of vGlut1 puncta devoid (grey) or associated to (red) mitochondria along the axon, and at branchpoints. N=22 axons out of 2 independent experiments. Bars: median ± 95% CI

We previously identified that presynaptic capture of mitochondria is required for axonal branching *in vitro* and *in vivo* (J. Courchet et al., 2013). However, since vGlut1-positive puncta outnumber mitochondria along the axon, it is unclear if all presynaptic boutons are equally important for axonal branching. We thus performed extensive correlation of mitochondria position and presynaptic boutons along the axon (**Fig. 2C**). As it is the case for mitochondria, the distance between vGlut1-positive puncta was lower than expected if puncta were evenly distributed, indicating that presynaptic boutons form clusters along the axon (**Fig. 2H**). These clusters were significantly closer and denser at branchpoints compared to along the axon (**Fig. 2I-J**). Virtually all branches had a vGlut1-positive cluster at their origin, and more than half of them also had resident mitochondria (**Fig. 2K**). Accordingly, presynaptic boutons located to branchpoint are more likely to be associated to mitochondria compared to boutons located along the axon (**Fig. 2L**). Finally, we tested if the distribution of synaptic puncta and mitochondria is conserved in an *in vivo* setting by performing *In Utero* Cortical Electroporation (IUCE) targeting callosal-projecting superficial (layer 2/3) PNs. The axon formed by these neurons projects toward the white matter (WM) and form collateral branches on layer 5 specifically (**Fig. S2A-B**). As expected, these axons form more synaptic puncta on layer 5 compared to the adjacent layers IV and VI. In comparison mitochondria density showed a progressive reduction with distance from the soma, but no specific enrichment in the branch-rich layer 5 (**Fig. S2C-D**). However, as we observed *in vitro*, the large majority of axonal branches had a vGlut1-positive cluster at their origin, and branch-associated vGlut1 puncta were more likely to harbor a mitochondrion than vGlut1 puncta along the axon (**Fig. S2E-F**). Taken together, our results strongly suggest that mitochondria are not randomly recruited to nascent presynaptic boutons but rather preferentially associate to vGlut1 puncta located at branch origin.

### Mitochondria are recruited after, and not prior to branch formation

Since our results presented so far relied primarily on static imaging, we next sought to determine the spatio-temporal dynamics of presynaptic bouton and mitochondria recruitment relative to branch formation. To do so we used EVCE and performed time-lapse imaging of nascent presynaptic boutons (defined as vGlut1 positive puncta) and mitochondria (marked with mito-DsRed) over 24 hours periods in 5-7DIV PNs (**Fig. S3A-B**). To alleviate phototoxicity while preserving spatio-temporal dynamics, we performed pulses of high-frequency imaging (5 minutes at 12 frames per minute) followed by periods with low frequency imaging (20 minutes at 0.1 frames per minute). We detected 45 events of spontaneous branch formation over 24 neurons and quantified presynaptic boutons and mitochondria localization at branchpoints and within branches, as well as axonal branch length and lifetime.

Studies in the Xenopus and Zebrafish demonstrated that axonal collateral branches from RGC neurons originate from presynaptic sites (Javaherian and Cline, 2005; Meyer and Smith, 2006; Ruthazer et al., 2006). We confirmed some of these findings in mouse cortical PNs (**Fig. S3C**): time-lapse longitudinal imaging revealed that presynaptic boutons form along the axon before the formation of collateral branches. Furthermore, these presynaptic boutons remained stable at branch origin and additional vGlut1 clusters formed inside the branch as it consolidates.

In mouse sensory axons, mitochondria recruitment at sites of local protein synthesis promote filopodia and branch formation (Spillane et al., 2013). However, in cortical PNs, we observed that mitochondria were recruited to branchpoints after, and not prior to, branch formation (**Fig. 3A** and **Fig. S3D**). Immediately following the turning of filopodia to new axonal branches (as defined by the formation of a growth-cone like structure, T=0min), mitochondria started clustering at branch origin. On average the number of mitochondria increased by 42% at branchpoints within 30 minutes of branch formation (**Fig. 3B**). In parallel we assessed local changes in mitochondria motility around axonal branch points (**Fig. 3C**). Following branch formation, we observed clustering of immobile mitochondria within 5µm of branch origin (green box in **Fig. 3C**). In contrast, mitochondria passing in the corresponding region of the axon before branch formation only displayed short dwelling periods. Individual mitochondria dwell time at branch origin was significantly longer than at other positions along the axon (**Fig. 3D**).

**Figure 3:**
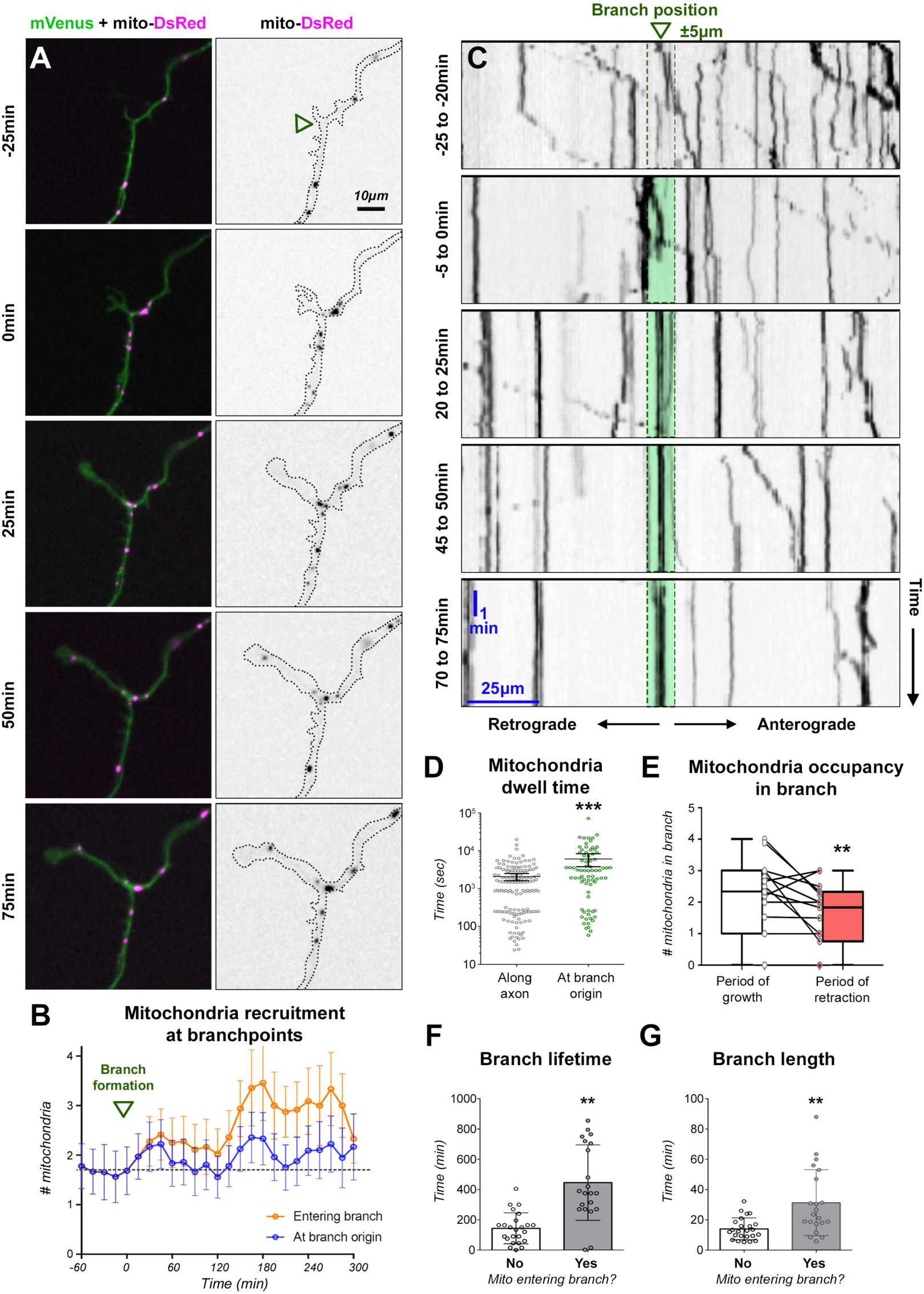
dynamics of mitochondria recruitment at branch origin and inside branches. (A) Time-lapse imaging of a spontaneous branch formation from the axon of a cortical PN at 5DIV following EVCE of mVenus (green) and mito-DsRed (magenta) coding plasmids. Branch apparition defined by length >5µm and a growth-cone like structure was considered T=0min. (B) Dynamics of mitochondria recruitment at branch origin (blue) or at branch origin and inside branch (orange) over time. Data: average ± SEM, N=12 axons. (C) Kymographs of axonal mitochondria centered around branch position (green arrowhead) from axon in (A). (D) Mitochondria dwell time at branch origin (± 5µm) compared to dwell time along the axon. Time was represented on a logarithmic scale. Each point represents a mitochondrion. N_(along axon)_=143, N_(branch origin)_=75 out of 14 axons. Bars: median ± 95% CI. Statistical test: Mann-Whitney. (E) Comparison of mitochondria occupancy in a branch during periods of growth (white) or retraction (red). Box-plot: 75^th^ percentile, median and 25^th^ percentile. Individual dots represent average over period (per branch), N=15 branches. Statistical test: Wilcoxon matched-pairs ranked test. (F-G) Correlation of mitochondria entry into a branch with lifetime (F) and length (G). Bars: median ± 95% CI. Each dot represents a spontaneous branch formation event. N_(No)_=24; N_(Yes)_=21. Statistical test: Mann-Whitney

Furthermore, we observed that mitochondria entered newly formed collateral branches shortly after clustering at branch origin (**Fig. 3A**, T=50min and **Fig. S3D**). Within 2 hours of branch formation, there was a marked increase of mitochondria entering branches (**Fig. 3B**). The number of mitochondria inside branches was correlated with periods of extension and decreased when branches were retracting (**Fig. 3E**). Accordingly, mitochondria entering branches were correlated with increased length and lifetime (**Fig. 3F-G**). On the contrary branches that failed to recruit mitochondria rapidly retracted (**Fig. S3E**). Overall, our results demonstrate that contrary to sensory axons, branch formation precedes mitochondria clustering but also revealed that mitochondria are involved in axonal branch stabilization in mouse cortical PNs.

### Mitochondria are required for branch stabilization

We next tested how the local pool of mitochondria is required for branch stabilization by performing Chromophore-Assisted Light Inactivation (CALI) of mitochondria (Bulina et al., 2006a; 2006b). As a proof of principle, we co-expressed the genetically-encoded photosensitizer Killer-Red (mito-KR) (Bulina et al., 2006b; 2006a) targeted to the mitochondrial matrix together with the biosensor mito-roGFP2 (Gutscher et al., 2008) to measure ROS production in mitochondria following photosensitization in HeLa cells. Following a single 30sec laser pulse, mito-KR induced a strong production of ROS in mitochondria restricted to the laser-targeted Region Of Interest (ROI) (**Fig. S4A-D**). Furthermore, CALI led to a rapid fragmentation of mitochondria in the ROI (**Fig. S4E-G**). We observed no change in roGFP2 signal in the cytosol even 30 minutes following CALI, suggesting minimal ROS leakage from damaged mitochondria (**Fig. S4H**). CALI on control fluorescent protein mt-DsRed did not induce significant morphological changes in mitochondria or accumulation of ROS (**Fig. S4D-G**).

We subsequently expressed mito-KR in cortical PNs by EVCE and targeted axonal mitochondria for photosensitization. As observed in HeLa cells, a 30sec laser pulse induced a rapid accumulation of ROS in mitochondria, restricted to the ROI (**Fig. 4A-B**). We then assessed the local metabolic consequences of axonal CALI by expressing PercevalHR, a genetically-encoded biosensor for the ATP:ADP ratio. Following laser illumination of mito-KR, we measured an immediate drop in PercevalHR signal, indicating a local decrease of the ATP:ADP ratio (**Fig. 4C**). The reduction in PercevalHR signal was persistent over 60 minutes and largely restricted to the ROI (**Fig. 4D-E**). In contrast PercevalHR signal was not affected by laser illumination upon expression of control mt-DsRed (**Fig. 4F**).

**Figure 4:**
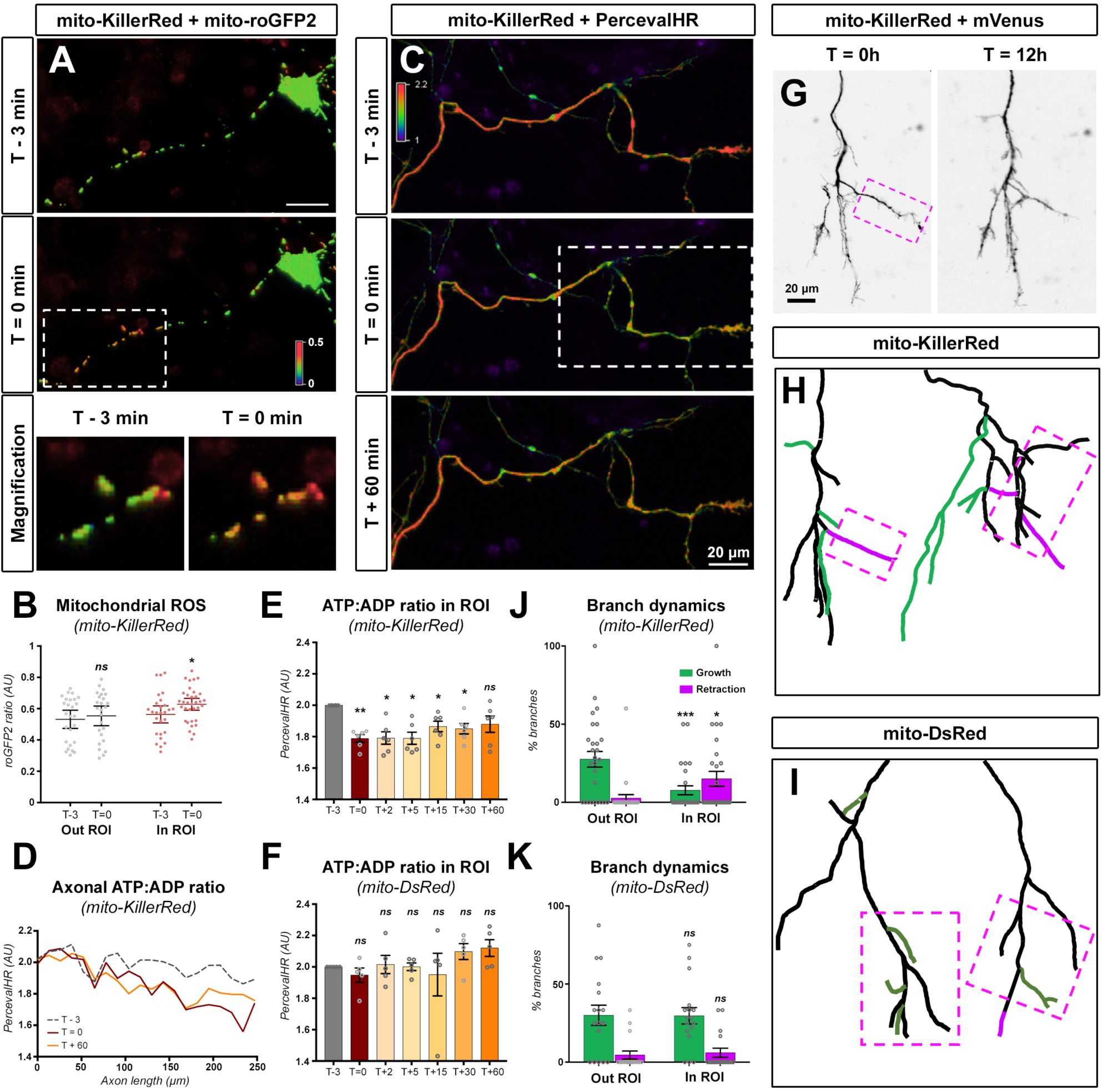
mitochondria CALI disrupts branch stabilization. (A) Typical neuron at 5DIV expressing mt-KR. At T=0min, CALI was performed by a 30sec laser pulse in the Region of Interest (ROI, white box). ROS was measured by mito-roGFP2 fluorescence (red = oxidized, blue = reduced). (B) Mitochondria ROS accumulation was measured outside and inside of the ROI before and immediately after CALI. Each point represents roGFP2 signal (Ex405:Ex488 ratio, see methods) for a given mitochondrion. Bars: average ± 95% CI. Analysis: Unpaired t-test. N_(T-3, OUT)_=26, N_(T=0, OUT)_=25, N_(T-3, IN)_=27, N_(T=0, IN)_=34 mitochondria out of 7 independent experiments. (C) Measurement of the ATP:ADP ratio with PercevalHR in the axon of cortical PNs expressing mt-KR. CALI was performed in the indicated ROI (white box). (D) PercevalHR signal (Ex488:Ex405 ratio, see methods) was measured along the axon in (C) at the indicated time-points. (E-F) Average PercevalHR signal in the ROI of neurons expressing mt-KR or the control mt-DsRed. Bars: average ± SEM. Analysis: Repeated measures ANOVA with Dunnett’s multiple comparison test. N=5 axons out of 5 independent experiments. (G) Time-lapse imaging of axon growth over 12 hours following CALI (magenta box). mVenus fluorescence was used to assess axonal morphology. (H-I) Examples of axon (schematic reconstruction) expressing mt-KR or the control mt-DsRed. Growing/new branches (green) and retracting branches (magenta) were observed over 12 hours and quantified in and out of the ROI (J-K). Bars: average ± SEM. Analysis: 2way ANOVA with Bonferroni’s multiple comparison test. Growth or retraction in ROI was compared to out ROI. N_KillerRed_=27 axons, N_DsRed_=17 axons out of 4 independent experiments.

Finally, we expressed mito-KR together with the green fluorescent protein mVenus by IUCE and observed axon morphogenesis over 12 hours time-lapse movies (**Fig. 4G**). Localized photoinactivation of mitochondria significantly altered branch dynamics in the ROI, by promoting branch retraction and decreasing branch growth, whereas this effect was not observed on the same axon outside of the ROI, indicating that overall axonal viability is not affected. In contrast, laser illumination had little effect on branch dynamics upon expression of mt-DsRed (**Fig. 4H-K**). Taken together, our results demonstrate that mitochondria are required locally to promote branch stabilization.

### NUAK1 kinase controls axonal mitochondrial metabolism

The observation that mitochondrial metabolism is necessary and sufficient to promote collateral branching (**Fig. 1**) and that mitochondria are required locally for axonal branch stabilization (**Fig. 3**) raises the question of their local function. We previously reported that the Serine/Threonine kinase NUAK1 controls terminal axon branching through the control of mitochondria capture at presynaptic sites in developing PNs (J. Courchet et al., 2013).

Interestingly NUAK1, a kinase related to the master metabolic regulator AMPK (Lizcano et al., 2004) is also involved in mitochondrial metabolism in c-Myc dependent cancer models (Liu et al., 2012). We therefore sought whether NUAK1 controls mitochondrial function as well in cortical PNs. Using a Seahorse analyzer, we measured the glycolytic and respiratory capacity of 7 DIV neuronal cultures from NUAK1^+/+^ (WT), NUAK1^+/−^ (HET) or NUAK1^−/−^ neurons (KO). There was a dose-response decrease in both the basal and maximal respiratory rate upon deletion of NUAK1, whereas glycolysis was largely unaffected (**Fig. 5A-B**). Importantly there was no difference in mitochondrial DNA content in NUAK1 KO neuronal cultures compared to WT conditions, thus ruling out that the decreased respiratory rate is a secondary consequence of impaired mitochondria biogenesis (**Fig. 5C**). To confirm these results, we assessed the overall metabolic consequence of NUAK1 knockout by measuring the ATP:ADP ratio with the biosensor PercevalHR. In 5DIV neuronal cultures following EVCE of plasmids encoding CRE together with PercevalHR, we observed a marked decrease of the ATP:ADP ratio in NUAK1 KO neurons (**Fig. 5D-E**). After quantification, the average PercevalHR signal was decreased by 21,9% in NUAK1 KO neurons compared to control cortical PNs (**Fig. 5F**). Interestingly, the decrease in PercevalHR signal was even more pronounced in the axon of NUAK1 KO neurons (−23.4%) (**Fig. 5G**).

**Figure 5:**
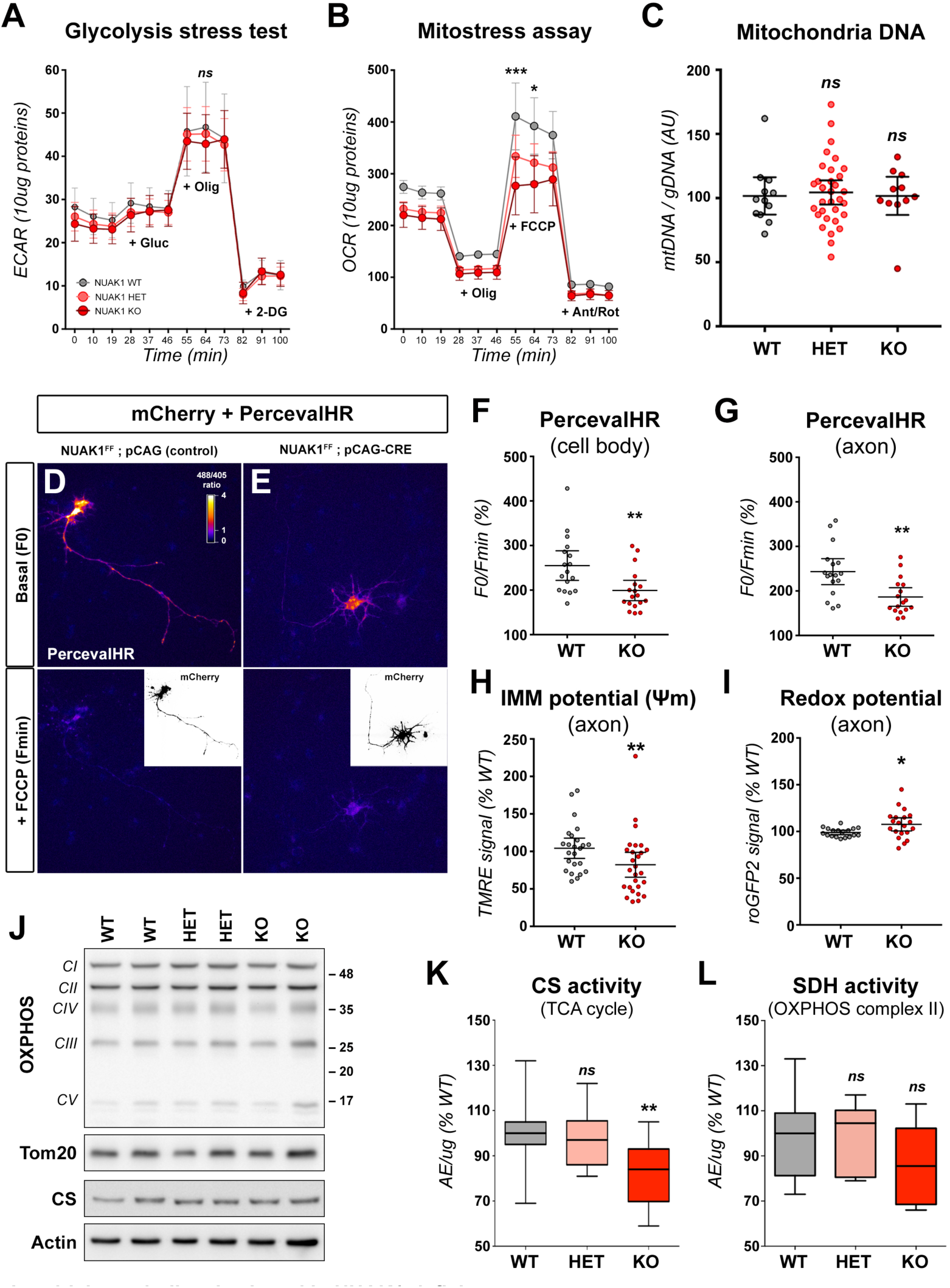
mitochondrial metabolism is altered in NUAK1-deficient neurons. (A-B) Measurement of glucose metabolism in 5DIV cortical neurons cultures from the indicated genotypes. ExtraCellular Acidification Rate (ECAR) and Oxygen Consumption Rate (OCR) were measured respectively to assess glycolysis and oxidative metabolism. Data: average ± SEM. N_WT_=9, N_HET_=27, N_KO_=9. (C) Measurement of mitochondrial content in cortical neurons cultures of the indicated genotypes. Mitochondrial and genomic DNA was measured by quantitative PCR. Each point represents a distinct culture. Bars: average ± 95% CI. Analysis: Kruskal-Wallis test with Dunn’s post-test. N_(WT)_=12, N_(HET)_=32, N_(KO)_=11. (D-E) Measurement of the ATP:ADP ratio in 5DIV NUAK1^F/F^ neurons transfected with a control (D) or a CRE coding plasmid (E). Excerpt: mCherry fluorescence was used to determine neuronal morphophogy. PercevalHR signal is displayed as a fluorescence ratio between 488 and 405 illumination using a Fire LUT (White: high ratio. Blue: low ratio) (F-G) Measurement of the ATP:ADP ratio in NUAK1^F/F^ neurons expressing a control (WT) or CRE-expressing plasmid (KO). Each point represents average values for a given neuron. Bars: average ± 95% CI. N_(WT)_=18, N_(KO)_=18 out of 3 independent experiments. (H-I) Measurement of the mitochondrial membrane potential (ΔΨm) or redox potential (GRX1-roGFP2) in the axon of NUAK1^F/F^ neurons expressing a control (WT) or CRE-expressing plasmid (KO). Each point represents average values for a given neuron. Bars: average ± 95% CI. Analysis: Unpaired t-test with Welch’s correction. (H) N_(WT)_=24, N_(KO)_=27 out of 5 independent experiments. (I) N_(WT)_=18, N_(KO)_=20 out of 4 independent experiments. (J) Typical Western-blot showing the expression of OXPHOS proteins in mitochondria from cortical neuronal cultures of the indicated genotypes. (K-L) SDH and CS enzymatic activity relative to protein quantity (normalized to WT). Box-plot: 75th percentile, median and 25th percentile. Analysis: One-way ANOVA with Dunnett’s multiple comparison test. N_(WT)_=13, N_(HET)_=8, N_(KO)_=10 independent samples.

The decreased respiratory rate suggested that NUAK1 is required not only for mitochondria trafficking, but also for proper mitochondria function in axons. To test this hypothesis, we turned to microscopy-based strategies to directly investigate mitochondrial function. Cortical neuron cultures were performed from floxed NUAK1 embryos (NUAK1^F/F^; (V. Courchet et al., 2018)) in order to achieve sparse, single-cell NUAK1 inactivation. CRE recombinase encoding plasmids were electroporated together with plasmids encoding the fluorescent protein mVenus and the mitochondrial marker mito-BFP. At 5DIV, cultures were exposed to Tetra-Methyl-Rhodamine Ethyl-ester (TMRE) to measure mitochondrial inner membrane potential (Ψ_m_). We observed a marked decrease of Ψ_m_ in the axon of NUAK1 KO neurons, compared to control cortical PNs electroporated with a non-CRE expressing plasmid (**Fig. 5H** and **Fig. S5A**). In parallel, we measured axonal mitochondrial redox potential using a mitochondria-targeted roGFP2. We observed a shift of roGFP2 signal toward a more oxidized state in NUAK1 KO neurons, suggesting an increased ROS production by axonal mitochondria (**Fig. 5I** and **Fig. S5B**).

To check whether the low Ψ_m_ and oxidized mitochondria state is a secondary consequence of the increased trafficking of NUAK1 KO neurons, we measured TMRE and roGFP2 fluorescence in wild-type neuronal cultures and sorted out motile versus stationary mitochondria (**Fig. S5C-F**), however there was no difference in either measured parameter.

Altogether, our results strongly argue that mitochondria function is decoupled from mitochondria trafficking in the axon and therefore suggest that NUAK1 regulates not only presynaptic mitochondria capture but also mitochondrial metabolic function. Finally, we turned to biochemical assays to determine if mitochondria are defective *per se* or if mitochondria impairment is context-dependent. Specifically we sought to confirm that the alterations in mitochondrial metabolism resulting from NUAK1 loss are conserved in an *in vivo* setting. We measured the activity of two key mitochondrial enzymes, Citrate Synthase (CS), a key enzyme of the Citric Acid Cycle, and of Succinate Dehydrogenase (SDH), forming the complex II of the mitochondrial respiratory chain. We observed a marked, dose-dependent reduction of CS from mitochondria extracted of the cortex of NUAK1 KO mice (**Fig. 5K**). We measured a similar, although not statistically significant, reduction in SDH activity from NUAK1 KO cortices (**Fig. 5L**). As a control, we checked the expression of OXPHOS supercomplexes and CS by Western-blot but observed no change with NUAK1 expression (**Fig. 5J**).

Taken together, our results indicate that NUAK1 controls not only mitochondria positioning but also mitochondria function along the axon and suggest that a metabolic imbalance may account for the axon branching phenotype of NUAK1 KO neurons.

### Upregulation of mitochondria function rescues collateral branching

In order to determine if there is a direct, causal, link between the downregulation of mitochondrial metabolic activity and the reduction in collateral branching in NUAK1 KO axons, we devised strategies to upregulate mitochondria function. Our prediction was that, improving mitochondrial fitness should rescue collateral branching in NUAK1 deficient neurons. Based on published studies, we treated cortical PNs with the cell-permeant mitochondria modulator L-Carnitine (Dickey and Strack, 2011; Sainath et al., 2017), a cell-permeant, lysine-derived quaternary amine that acts by promoting the metabolism of long-chain fatty acids by the mitochondria. It has been shown to be neuroprotective in a wide variety of metabolic stresses (Tefera and Borges, 2016; Zanelli et al., 2005) and has clinical interest in the treatment of depression and anxiety (Chiechio et al., 2017; Filiou and Sandi, 2019).

We achieved single-cell knockout of NUAK1 through EVCE of CRE-coding plasmid in NUAK1^F/F^ cortical PNs and quantified axon morphology at 5DIV. As previously reported (J. Courchet et al., 2013) NUAK1-null neurons had shorter axons (**Fig. 6A**) with a reduced number of collateral branches (red stars in **Fig. 6A**). Upon L-Carnitine addition to the culture medium, we observed no effects on axon length in control or NUAK1-null cortical PNs (**Fig. 6B**). In contrast, L-Carnitine addition to NUAK1-deficient neurons (but not control) increased collateral branch formation compared to untreated control or NUAK1-deficient neurons. When normalized to axonal length, L-Carnitine induced a complete rescue of collateral branches, without affecting axon elongation (**Fig. 6B-C**). Importantly, L-Carnitine treatment had no effect on wild-type neurons. As an alternative means to upregulate mitochondrial function, we treated neuronal cultures with the mitochondria-targeted electron carrier Vitamin K2/menaquinone (VK2) (Vos et al., 2012). Similar to the effect of L-Carnitine, we observed a normalization of axonal branching in NUAK1-deficient neurons without affecting axonal length in VK2 treated neurons (**Fig. S6A-B**).

**Figure 6:**
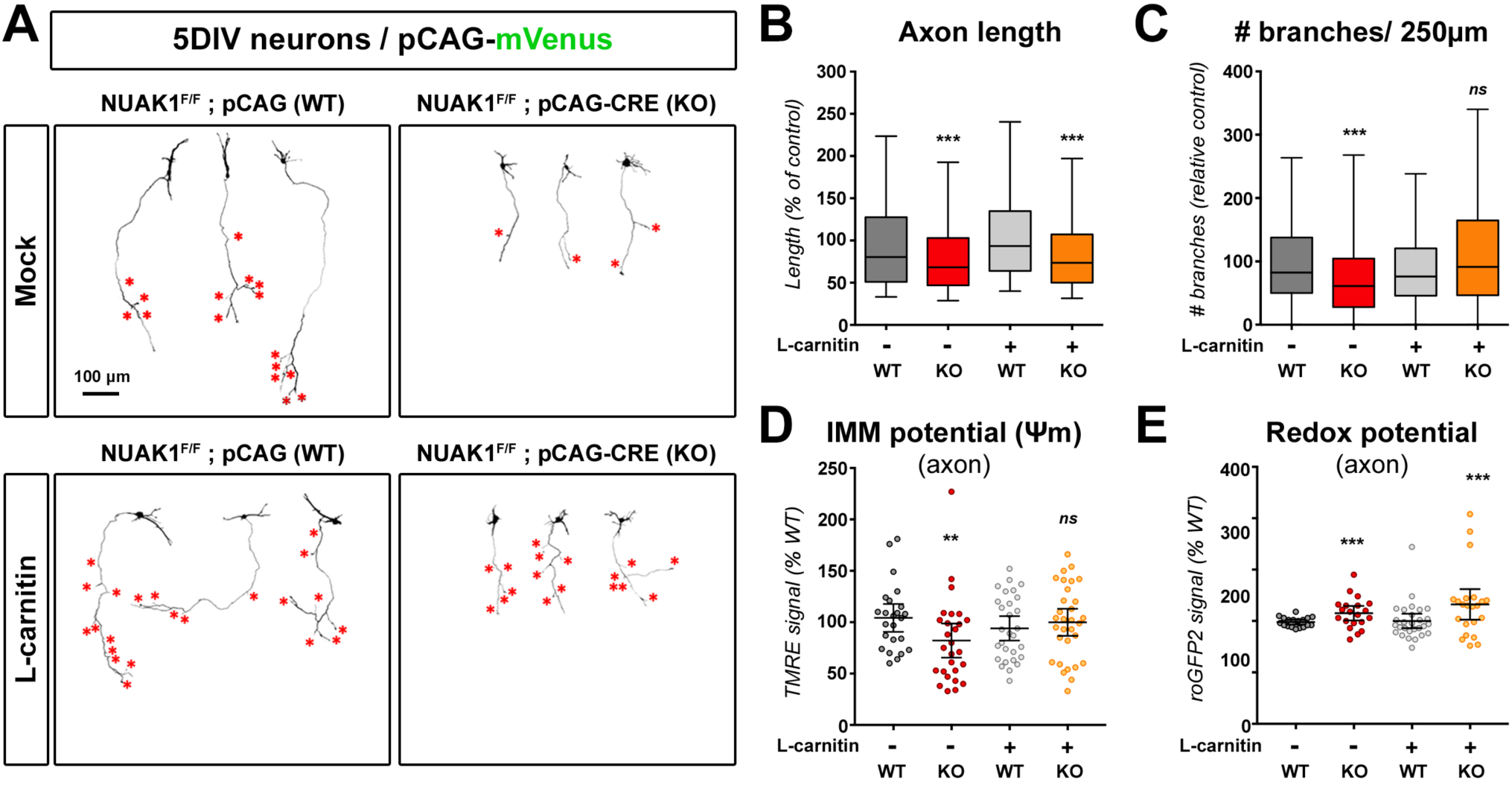
upregulation of mitochondrial function rescues axon branching in NUAK1-deficient neurons. (A) Representative images of mVenus expressing cortical neurons of NUAK1F/F background and electroporated with a control or CRE-coding plasmid. At DIV2 neurons were treated with L-Carnitine (1mM). Red stars indicate collateral branches. (B-C) Quantification of axon length and collateral branches of 5DIV neurons in the indicated conditions. Box-plot: 75^th^ percentile, median and 25^th^ percentile. Statistical tests: Kruskal-Wallis test with Dunn’s post-test (each condition compared to untreated WT condition). N_(WT)_=399, N_(KO)_=232, N_(WT+L-Carn)_=387, N_(KO+L-Carn)_=256 out of 5 independent experiments. (D-E) Effect of L-Carnitine on the mitochondrial membrane potential (ΔΨm) and redox potential of WT and KO neurons. Untreated samples from experiment in Figure 5D-E. Each point represents average values for a given neuron. Bars: average ± 95% CI. Analysis: Unpaired t-test with Welch’s correction. (D) N_(WT+L-Carn)_=29, N_(KO+L-Carn)_=31 out of 5 independent experiments. (E) N_(WT+L-Carn)_=28, N_(KO+L-Carn)_=22 out of 4 independent experiments.

We next sought to confirm that L-Carnitine treatment upregulated mitochondrial function in NUAK1-null neurons. We turned to microscopy-based assays using TMRE to measure Ψ_m_ in 5DIV neuronal cultures. L-Carnitine treatment normalized Ψ_m_ in NUAK1 KO neurons (**Fig. 6D**), indicating an upregulation of mitochondrial fitness. Interestingly, L-Carnitine had no effect on mitochondrial redox potential and failed to rescue the increased mito-roGFP2 signal in NUAK1-null neurons (**Fig. 6E**), thus ruling out that L-Carnitine rescues axonal branching through ROS scavenging. Accordingly, application of the ROS scavenger N-AcetylCystein (N-AC) had no effect on axon branching despite the normalization of oxidative stress in treated neurons (**Fig. S6C-E**).

### L-Carnitine rescues terminal axon branching in NUAK1 KO *in vivo*

Finally, we performed experiments to determine if metabolic upregulation can rescue terminal axon branching in NUAK1 KO mice *in vivo*. Our choice turned toward L-Carnitine because it presented several advantages: first it is virtually not toxic and especially it is used in humans for treatment in newborn babies to treat metabolic disorders and carnitine deficiency. Second, it is suitable for oral administration, and has been approved for use in veterinary medicine. We performed a conditional inactivation of NUAK1 in the dorsal telencephalon by breeding with the Nex^CRE^ mouse line (Goebbels et al., 2006) to induce cortex-specific post-mitotic deletion in all cortical PNs. We previously demonstrated that this conditional genomic deletion leads to a significant decrease in terminal axon branching of contralaterally-projecting layer 2/3 axons (V. Courchet et al., 2018). Upon detection of gestation (at E13.5), time pregnant mice were treated with L-Carnitine in the drinking water, while the control group received only regular water. At E15.5, we performed *In Utero* Cortical Electroporation (IUCE) of plasmid coding the red fluorescent protein mScarlet-I as a neuronal cell filler (Bindels et al., 2017). Drug treatment was continued after birth and during lactation, and pups were sacrificed at weaning (P21) when cortical layer 2/3 PNs axons display adult-like branching patterns (**Fig. 7**).

**Figure 7:**
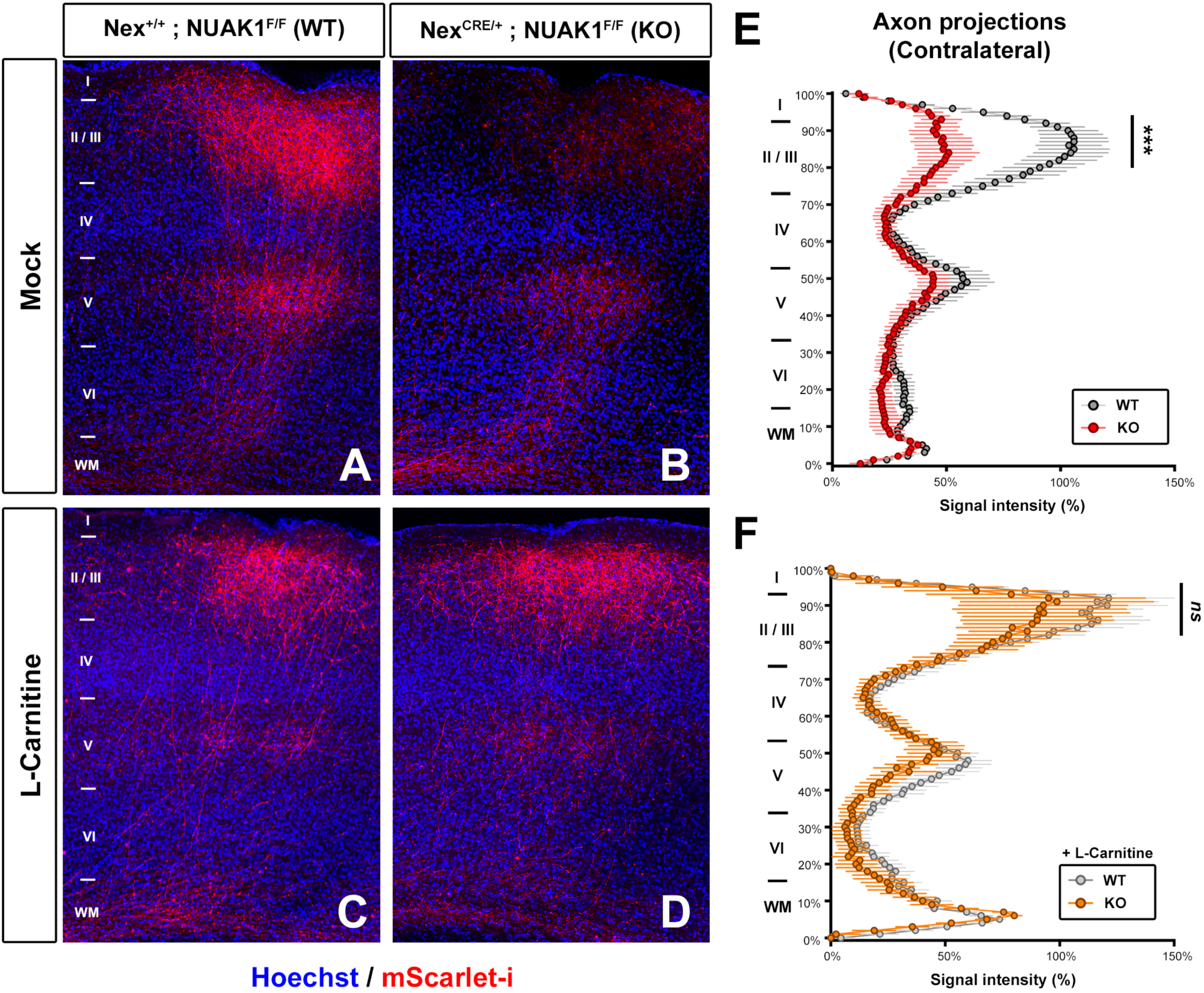
L-Carnitine treatment rescues terminal axon branching *in vivo*. (A-D) Representative images of contralateral terminal axon branching in NexCRE; NUAK1 mice following IUCE with a plasmid coding the red fluorescent protein mScarlet-I. When indicated (C-D) animals were treated with L-Carnitine in drinking water. The observed decrease in terminal axon branching in KO animals (B) was rescued upon L-Carnitine treatment (D). (E-F) Quantification of normalized mScarlet-i fluorescence along a radial axis in the contralateral cortex. Data: Average ± SEM. Statistical tests: Two-way ANOVA with Bonferroni’s multiple comparison. N_WT_=8, N_KO_=6, N_WT+L-CAR_=10, N_KO+L-CAR_=6

As expected, we could observe a decrease in terminal axon branching in KO mice (Nex^CRE/+^;NUAK1^F/F^) compared to controls (Nex^+/+^;NUAK1^F/F^) (**Fig. 7A-B**). On the contrary, terminal axon branching was undistinguishable between WT and KO mice upon L-Carnitine treatment (**Fig. 7C-D**). Quantification confirmed that L-Carnitine treatment during pregnancy and lactation is sufficient to rescue axon branching, especially in contralateral layer 2/3 (**Fig. 7E-F**). Taken together, our results suggest that metabolic upregulation of mitochondrial function can rescue defective axon branching in NUAK1 deficient mice *in vivo*.

## DISCUSSION

In the present study, we extended our previous observations that mitochondria distribution in the axon is necessary and sufficient for terminal branching, by providing new evidence that mitochondria positioning at axonal branch points and mitochondria function locally supports branch stabilization. We characterized that mitochondria capture at presynaptic boutons occurs preferentially at boutons associated with axonal branch points. Finally, we observed that the AMPK-related kinase NUAK1 controls not only axonal mitochondria distribution but also mitochondrial function and that upregulation of mitochondrial metabolism is sufficient to rescue the axonal branching defects characterizing NUAK1-deficient axons *in vitro* and *in vivo*. Together our results suggest a two-hit model by which it is equally important to ensure that mitochondria reach their proper destination, but also that they are active locally, to stabilize nascent axonal branches.

One striking observation from our study is the fact that axon elongation and collateral branching are operatively decorrelated in terms of metabolic requirement. First, we observed a dose-response effect of D-glucose concentration on axon length, whereas collaterals formation could be supported in an all-or-nothing fashion by directly fueling mitochondrial metabolism (**Fig 1**). Furthermore L-Carnitine and VK2 can rescue axon branching in a NUAK1 null background, without affecting significantly axon length (**Fig 6**). We previously reported that an autism-linked truncated NUAK1 impairs axon branching without affecting axon elongation (V. Courchet et al., 2018), although mitochondrial function has not been explored for this mutant. Taken together these observations suggest that, in cortical axons, mitochondrial function is necessary and sufficient for collateral branching. Further studies are warranted to determine if axon elongation involves distinct metabolic pathways, and if so whether these are also dependent upon NUAK1 function.

Several lines of evidence support the idea that immature neurons differ significantly from synaptically active neurons in terms of their metabolic organization. Axonal transport of mitochondria plays a critical role in supporting axon morphogenesis (Devine and Kittler, 2018; Misgeld and T. L. Schwarz, 2017), yet mitochondria motility decreases gradually as neurons mature (Lewis et al., 2016; Moutaux et al., 2018; Obashi and Okabe, 2013). Specifically, mitochondria are recruited at immature presynaptic boutons (J. Courchet et al., 2013) where they remain captured by an actin-dependent mechanism (Gutnick et al., 2019). Here, we further demonstrate that mitochondria are recruited after, and not prior to, branch formation. Furthermore, mitochondria are not recruited to any given presynaptic boutons, but are preferentially recruited to branch-associated boutons. This differs from observations in sensory and retinal neurons where mitochondria recruitment to hot-spots of local translation induce filopodia and branch formation (Spillane et al., 2013; Wong et al., 2017). It is possible that this discrepancy stems from distinct modalities of collateral branch formation. Indeed cortical PN branches emerge from regions where the growth cone paused during axon elongation (Kalil and Dent, 2014; Szebenyi et al., 1998), whereas NGF-induced branch can form anyplace along the axon (Gallo and Letourneau, 1998). Interestingly although NUAK1 expression is largely enriched in the developing cortex and hippocampus (Hirano et al., 2006), studies suggest other AMPK-RKs can functionally operate in regulating axon branching in sensory neurons (Lilley et al., 2013), potentially explaining mechanistic differences in the modalities of branch formation in different neuron types. Of note our study does not rule out a possible role of NUAK1 and mitochondria recruitment in supporting local protein synthesis in cortical neurons.

Our results reveal that mitochondrial function is needed to support axonal branching in connection with presynaptic boutons. Mature synapses have high metabolic need. Recent studies identified that synaptic ATP comes mostly from glucose uptake and metabolism locally (Ashrafi et al., 2017) and involves the two main components of glucose, ie. glycolysis (Jang et al., 2016; Rangaraju et al., 2014) and mitochondrial oxidative phosphorylations (Ashrafi et al., 2020; Pathak et al., 2015). However, it is unlikely that ATP by itself is sufficient to support branch formation and stabilization. Indeed the metabolic functions of mitochondria are not restricted to ATP production but contribute to most metabolic pathways in the cell including the metabolism of lipids, amino acids, nucleic acids or neurotransmitters (Martínez-Reyes and Chandel, 2020). Furthermore, axon outgrowth and branching is dependent on local calcium dynamics (Hutchins and Kalil, 2008) and calcium buffering by mitochondria is important for cortical PN axons branching (Lewis et al., 2018). Mitochondrial ATP production and calcium dynamics are intertwined (Ashrafi et al., 2020; Llorente-Folch et al., 2013). Further studies are warranted to carefully dissect the metabolome of mitochondria captured at axonal branchpoints in developing axons.

Although our evidence clearly indicate that mitochondrial function is impaired in NUAK1 deficient neurons, the molecular link between NUAK1 and mitochondria is so far unknown. In c-Myc dependent cancer cells, a knockdown of NUAK1 downregulates the expression of several essential mitochondrial proteins including multiple components of the Electron Transport Chain (ETC) (Liu et al., 2012). This broad effect on mitochondrial proteome suggests that the kinase NUAK1 might have an indirect effect on mitochondrial composition. Interestingly NUAK1 has recently been linked to RNA biogenesis (Cossa et al., 2020). Furthermore NUAK1 is an AMPK-related kinase (Lizcano et al., 2004), and AMPK activity is decreased upon NUAK1 knockdown in cancer cells (Liu et al., 2012). Interestingly AMPK acts on mitochondria biogenesis through the transcription factor PGC1a, and on mitochondria dynamics through the Mitochondrial Fission Factor MFF, which are both involved in axon morphogenesis (Lewis et al., 2018; Vaarmann et al., 2016). The potential role of a NUAK1/AMPK crosstalk on axonal mitochondria function and axonal branching remains to be studied. Finally, NUAK1 has a well-characterized action on actomyosin organization through the phosphorylation of the myosin phosphatase regulator MYPT1 (Zagórska et al., 2010). Interestingly a recent study uncovered that actomyosin contractility promotes axonal transport, and specifically that actomyosin affects the speed of mitochondrial transport in the axon (T. Wang et al., 2020). Furthermore in sensory neurons actomyosin contraction at branchpoint antagonizes the formation of microtubule bundles in NGF-induced branch formation (Ketschek et al., 2015). To which extent a NUAK1/MYPT1 complex controls mitochondria positioning and axon morphogenesis through the regulation of actomyosin should be explored.

Mitochondrial defects can cause an increased ROS production, which has been linked to defects in cortical circuits formation (Fernandez et al., 2019). The use of ROS scavenger can alleviate some of the anatomical and behavioral phenotypes in a mouse model of schizophrenia (Cabungcal et al., 2014). However, we did not observe any benefit of N-AC treatment in NUAK1-deficient neurons. On the contrary L-Carnitine rescued axonal branching *in vitro* and *in vivo*. L-Carnitine has demonstrated some beneficial effects on axonal development and branching in sensory neurons (Sainath et al., 2017). Furthermore L-Carnitine has anti-depressant effect mediated by an upregulation of metabolic activity (Cherix et al., 2020; Filiou and Sandi, 2019). Our work provides further evidence that L-Carnitine might be of therapeutic interest in correcting mitochondrial disfunction and metabolic imbalance during neurodevelopment.

## Supporting information

Resources table

## ACKNOWLEDGEMENTS

The authors thank members of the Courchet lab and Institut NeuroMyoGène for useful comments and discussion. We are sincerely grateful to Franck Polleux for his support in building this project. We wish to thank Franck Polleux, Tommy Lewis, Yusuke Hirabayashi, SeokKyu Kwon, Rémi Mounier, Hélène Puccio and Thomas Boulin for technical advice, suggestions, and critical reading of the manuscript, and Agnès Duplany, Anne Devin and Arnaud Mourier for suggestions and technical help in setting up biochemical assays. We thank the personnel from the SCAR and ALECS-SPF mouse facility for animal care. This work was supported by the Fondation pour la Recherche Medicale (AJE20141031276) and ERC Starting Grant (678302-NEUROMET). M.L. was the recipient of a grant from AFM-telethon.

M.L. and J.C. conceived the experiments and interpreted the results. M.L., R.D.R., C.B., L.J. and A.A. performed the experiments. J.C. and G.M-D. performed and interpreted IUCE experiments. M.L. and A.G. performed neuronal cultures. J.C. prepared the manuscript.

## MATERIALS AND METHODS

### Animals

Mice breeding and handling was performed by following National Institutes of Health guidelines and the French and European legislation. Experimental protocols were approved by the CECCAPP Ethics committee (C2EA15) of the University of Lyon. Time-pregnant females were maintained in a 12hr light/dark cycle and obtained by overnight breeding with males of the same strain. Noon following breeding was considered as E0.5. Floxed NUAK1 mice (Nuak1^tm1a(KOMP)Wtsi^) have been described previously (V. Courchet et al., 2018). The NexCRE mouse line (Neurod6^tm1(cre)Kan^) (Goebbels et al., 2006) were provided by Sandrine Humbert (Grenoble Institute of Neuroscience, France). Animals were maintained on a C57Bl/6J background.

### DNA and plasmids

Endotoxin-free plasmid DNA was obtained using Macherey Nagel midi-prep kit. We used the following, previously described plasmids: empty vector pCAG-IRES-GFP (pCIG2), CRE-expressing vector pCAG-CRE-IRES-GFP (pCIG2-CRE) (Hand et al., 2005), mVENUS expressing vector pSCV2 (Hand and Polleux, 2011), pLKO, pCIG2-shSNPH, pCAG-mitoDsRed (J. Courchet et al., 2013), pFUGW-PercevalHR (Tantama et al., 2013), pCAG-mTagBFP2 and pCAG-mito-mTagBFP2 (Lewis et al., 2016).

pCAG-mito-KillerRed was created by cloning the photosensitizer KillerRed targeted to the mitochondria with two mitochondria targeting sequences (MTS, derived from subunit 8 of human cytochrome C oxidase) into a pCAG vector between XhoI and NotI sites, following PCR amplification from a pCMV-mito-KillerRed plasmid (Evrogen). mScarlet-i cDNA was amplified by PCR from a pmScarlet-i_C1 plasmid (gift from Dorus Gadella, Addgene #85044) (Bindels et al., 2017) and cloned into a pCAG2 backbone between XhoI and EcoRI sites. pCAG-mito-Grx1-roGFP2 and pCAG-cyto-Grx1-roGFP2 (Gutscher et al., 2008), pCAG-CRE, and pCAG-vGlut1-GFP (Herzog et al., 2011) were a kind gift from Tommy L Lewis (OMRF, USA), Seok-Kyu Kwon (KIST, Korea) and Etienne Herzog (U Bordeaux, France), respectively.

### Primary neuronal culture

The electroporation of dorsal telencephalic progenitors was performed by injecting plasmid DNA (1-2 μg/μL of endotoxin-free plasmid DNA) plus 0.5% Fast Green (Sigma; 1:20 ratio) using a Picospritzer III microinjector (Harvard Apparatus) into the lateral ventricles of isolated E15.5 embryonic mouse heads (Polleux and Ghosh, 2002). Electroporation were performed with gold-coated electrodes (GenePads 5 mm, BTX) using an ECM 830 electroporator (BTX) and the following parameters: five pulses of 100 msec, 150 msec interval, at 20 V. Immediately after electroporation, cortices were dissected in Hank’s buffered salt solution (HBSS) supplemented with HEPES (pH 7.4; 2.5 mM), CaCl2 (1 mM, Sigma), MgSO4 (1 mM, Sigma), NaHCO3 (4 mM, Sigma), and D-glucose (30 mM, Sigma), hereafter referred to as complete HBSS (cHBSS). Isolated cortices were dissociated in cHBSS containing papain (Worthington, 20U/mg at least) for 20 min at 37°C. Cortices were washed once in cHBSS containing DNase I (2.5mg/mL, Sigma), then 3 times in cHBSS before being manipulated by trituration. Cells were then plated at 125.10^3^ cells per 35 mm glass bottom dish (MatTek) coated with poly-D-lysine (0.1mg/mL) and laminin (0.01mg/mL) and cultured for 5–7 days in Neurobasal medium supplemented with B27 (1x), N2 (1x), Glutamax (2mM), and penicillin (10U/mL)-streptomycin (0.1mg/mL).

### *In Utero* Cortical Electroporation

Timed-pregnant NUAK1^F/F^ females were mated with Nex^CRE/+^; NUAK1^F/F^ males. At E15.5, we performed in Utero Cortical Electroporation as described in (Meyer-Dilhet and J. Courchet, 2020). A mix containing 1 μg/μl endotoxin-free plasmid DNA (pCAG-mScarlet-i) plus 0.5% Fast Green (Sigma; 1:20 ratio) was injected into one lateral hemisphere. Electroporation was performed using an ECM 830 electroporator (BTX) using four pulses of 45V with 500 mSec interval to target cortical progenitors. Animals were sacrificed at Postnatal day 21 by terminal perfusion of 4% paraformaldehyde (PFA, Electron Microscopy Sciences) followed by 2 h post-fixation in 4% PFA. In the treated group, L-Carnitine (Isulik 20, Sogeval laboratories) was provided in the drinking water from detection of pregnancy (E13.5) to P21.

### Drug treatments

Unless indicated otherwise, drugs were from Sigma-Aldrich and diluted in water (see Supplementary Resource Table). For experiments of mitochondria inactivation (Figure 1), cells were treated with low doses or Rotenone (10, 25 or 50nM). For *in vitro* rescue experiments (Figure 6), neurons were treated at 2 DIV with either L-Carnitine (1mM), Vitamin K2 (Menaquinone K4) (1µM in ethanol), N-Acetyl-L-Cystein (1mM).

For measurements of mitochondrial Inner Membrane potential, neurons were pre-incubated in cHBSS during 30 min at 37°C, then incubated in cHBSS containing 20nM of TMRE during 30min at 37°C. Medium was replaced by cHBSS containing 5nM of TMRE immediately before live-imaging.

### Immunostaining histochemistry

Cells were fixed for 15 min at room temperature in 4% (w/v) paraformaldehyde in PBS, washed 3 times in PBS, then permeabilized for 1 hr in Permeabilization buffer (PB: 0.3% Triton X-100, 0.3% BSA (Sigma), in PBS). Primary antibodies were incubated for 2 hr at room temperature in PB. Secondary antibodies were incubated for 1 hr at room temperature. Coverslips (BioCoat) were mounted on slides with Fluoromount G (EMS). Primary antibodies used for immunohistochemistry and immunocytochemistry in this study are chicken anti-GFP (Rockland) (1:2000) and anti-CRE (Millipore) (1:1000). All secondary antibodies were Alexa-conjugated (Invitrogen) and used at a 1:1000 dilution. Nuclear DNA was stained using Hoechst 33258 (1:10,000, Pierce).

### Immunohistochemistry

At P21, mice were sacrificed by intracardiac perfusion of PFA (4% in PBS) followed by a 2 hours post-fixation of the whole brain in a 4% PFA solution. 75µm thick sections were performed using a Leica VT1000S vibratome. Slices were permeabilized for 30 minutes in PB, then incubated overnight with primary antibodies (chicken anti-GFP, Rockland) diluted at 1:2000 in PB. The following day, we performed 3 washes in PBS 1X, then incubated slices in secondary antibody-containing PB (Goat anti-chicken antibody, Alexa 488, 1:2000, life technology). Nuclear DNA was stained using Hoechst 33258 (1:5000).

### Western blotting

Neurons were lysed in ice-cold Lysis Buffer (LB) containing 50mM Tris (ph8) (Euromedex), 0,5% sodium deoxycholate (Sigma), 0,1% SDS (Sigma), 600 mMNaCl (Sigma) and 1X protease and 1X phosphatase mixture inhibitors (Roche). Neurons put in Bioruptor (Diagenode) for break nucleus and DNA (8 cycles with 15sec in Time on and 60sec in time off). 25μg aliquots of lysate were separated by electrophoresis on a 12% and 15% SDS-polyacrylamide gel and transferred on a polyvinylidene difluoride (PVDF) membrane (Amersham). Incubation with primary antibody was performed overnight in TBS, 0,1% Tween, 5% BSA and anti-OXPHOS antibody (abcam 110413) to 1/1000 dilution or anti-Actin antibody (MP Biomedicals) to 1/5000 dilution or anti-TOM20(Cell Signaling) to 1/1000 dilution. The next day, membranes were incubated at room temperature for 1 hr with HRP-linked secondary antibodies in TBS, 0,1% Tween. Western-blot revelation was performed using ECL Prime (Merk, Sigma). Imaging and analysis were performed on a chemidoc imager (Bio-rad) using the ImageQuant software. Images presented in the figures have been cropped around the expected bands for figure design purpose.

### Metabolomic flux analyzes

We used a Seahorse XFe24 analyzer (Agilent) with the Glycolysis and Mito Stress test kits. Neurons were plated at 100,000 cells per well in Seahorse compatible 24 well plates and metabolic activity was measured at 7 days *in vitro*. The assay was conducted in Modified HBSS (1XHBSS supplemented with glucose (3.5g/L), Sodium pyruvate 1mM, Glutamax 1X, CaCl2 1mM, MgSO4 1mM, pH 7.4). Following measurement, neurons were lysed in ice-cold lysis buffer (LB) and protein concentration was measured by Bradford for post-hoc normalization.

### Mitochondria DNA

Mitochondrial DNA was measured by qPCR following a protocol described elsewhere (Oruganty-Das et al., 2012). 7DIV primary neurons DNA was extracted by whole cell lysis in lysis buffer (Tris pH8 100mM, EDTA 5mM, SDS 0.2%, NaCl 200mM, proteinase K 0.5mg/mL) followed by isopropanol precipitation. Quantitative PCR was performed with the CFX-connect thermal cycler (BioRad). The following primers were used: mtDNA_F: ACCATTTGCAGACGCCATAA, mtDNA_R: TGAAATTGTTTGGGCTACGG, betatub_F: GCCAGAGTGGTGCAGGAAA, betatub_R: TCACCACGTCCAGGACAG.

### Biochemical Assay

Biochemical assays were performed on mouse cortex samples from NUAK1 KO, NUAK1 HET or NUAK1 WT mice. Samples were subjected to a chemical lysis using RIPA buffer (150mM NaCl, 0,5M EDTA ph8, 50mM Tris pH 8; 1% NP-40, 0.5% sodium deoxycolate, 0.1% SDS) and followed by a mechanical lysis using the Precellys Evolution.

Enzymatic activity (EA) was determined by measuring absorbance using a PowerWave XS plate reader (BioTek). We used the Beer Lambert law EA= Mean V(ε x l) where l is the width of the tank (l = 0.4 cm) and Mean V was expressed in mDO/min. For Succinate deshydrogenase activity assay, ε = 21000 uA/M/cm. For Citrate synthase activity assay, ε = 13600 uA/M/cm.

SDH activity was measured from a volume of 10µL (50µg of total proteins) diluted in 95µL of specific reaction buffer (65µL of 50mM KH_2_PO_4_, pH 7.2, 15µL of 70mM Succinate, 15µL of 560µM DCIP). Absorbance at 412nm was performed over 15 minutes at 37°C (one read per 10 seconds).

CS activity was measured from a volume of 20µL (100µg of total proteins) diluted in 90µL of specific reaction buffer (65µL of 50mM KH_2_PO_4_, pH 7.5, 10µL of Acetyl-CoA 4.4mM, 10µL of Oxaloacetate 4.4mM, 10µL of 150mM DTNB). Absorbance at 600nm was performed over 15 minutes at room temperature (one read per 10 seconds).

### Image Acquisition and Analyses

Confocal images were acquired in 1024×1024 mode with a Nikon Ti-E microscope equipped with the C2 laser scanning confocal microscope. Time-lapse images were acquired in 1024×1024 mode using a CMOS ORCA-Flash 4.0 Camera (Hamamatsu). Microscope control and image analysis was performed using the Nikon software NIS-Elements (Nikon). We used the following objective lenses (Nikon): 10x PlanApo; NA 0.45, 20x PlanApo VC; NA 0.75, 60x Apo; NA 1.4. When live-cell imaging was carried out, the temperature was maintained at 37° C and CO2 at 5% v/v in a humidified atmosphere. Live-cell imaging was performed in cHBSS. Representative neurons were isolated from the rest of the image using ImageJ. Contrast was enhanced and background (autofluorescence of no transfected neurons in culture) removed for better illustration of axons morphology. Kymographs were created with NIS-Elements.

### Quantifications and statistical analyses

Statistical analyses were performed using Prism (GraphPad). Statistical tests and number of replicates are indicated in figures legend. For axonal morphogenesis experiments, we performed large field acquisition of neuronal cultures (typically a 4×4 tiling scan using a 20x objective). All neurons on the reconstituted image were quantified and axon length was measured with the Nikon NIS Elements software. Axons shorter than 80µm were not counted. Quantifications were performed blind to genotype.

