## Supplementary figures 1-6 for "The AMPK-related kinase NUAK1 controls cortical axons branching though a local modulation of mitochondrial metabolic functions"

### Supplementary material

### SUPPLEMENTARY FIGURES

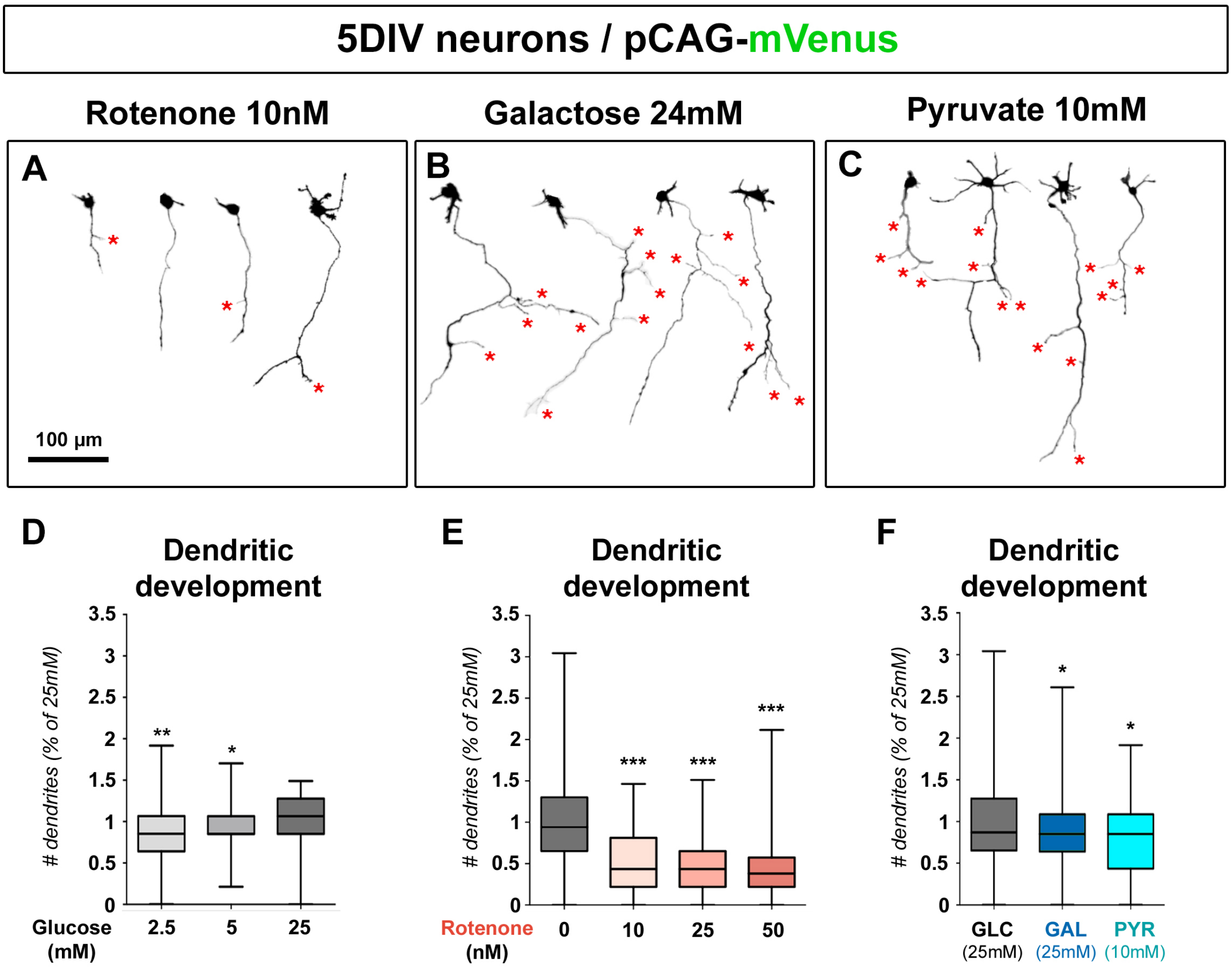


**Supplementary Figure 1 (related to Figure 1): the nature of the metabolic fuel in the medium has little impact on dendritic development at 5DIV**

(A) Representative images of mVenus expressing cortical neurons (5DIV) cultured in Neurobasal medium containing 25mM of glucose and treated with the indicated dose of Rotenone.

(B-C) Representative cortical neurons (5DIV) cultured in Neurobasal medium containing 24mM of galactose + 1mM of glucose (B), or cultured in Neurobasal medium containing 10mM of pyruvate without glucose (C).

(D-F) Quantification of the number of dendrites of 5DIV neurons in the indicated conditions (normalized to control 25mM condition): (D) medium containing increasing concentrations of glucose. (E) 25mM glucose-containing medium with increasing doses of rotenone. (F) medium containing either glucose (25mM), or galactose (24mM) + glucose (1mM), or Pyruvate (10mM) without glucose.

Box-plot: $75^{th}$ percentile, median and $25^{th}$ percentile. Statistical tests: Kruskal-Wallis test with Dunn's post-test (each condition compared to untreated 25mM condition). (D) $N_{(2.5mM)}$=148, $N_{(5mM)}$=198, $N_{(25mM)}$=99. (E) $N_{(0nM)}$=142, $N_{(10nM)}$=185, $N_{(20nM)}$=137, $N_{(50nM)}$=100. (F) $N_{(Glucose)}$=138, $N_{(Galactose)}$=210, $N_{(Pyruvate)}$=124.

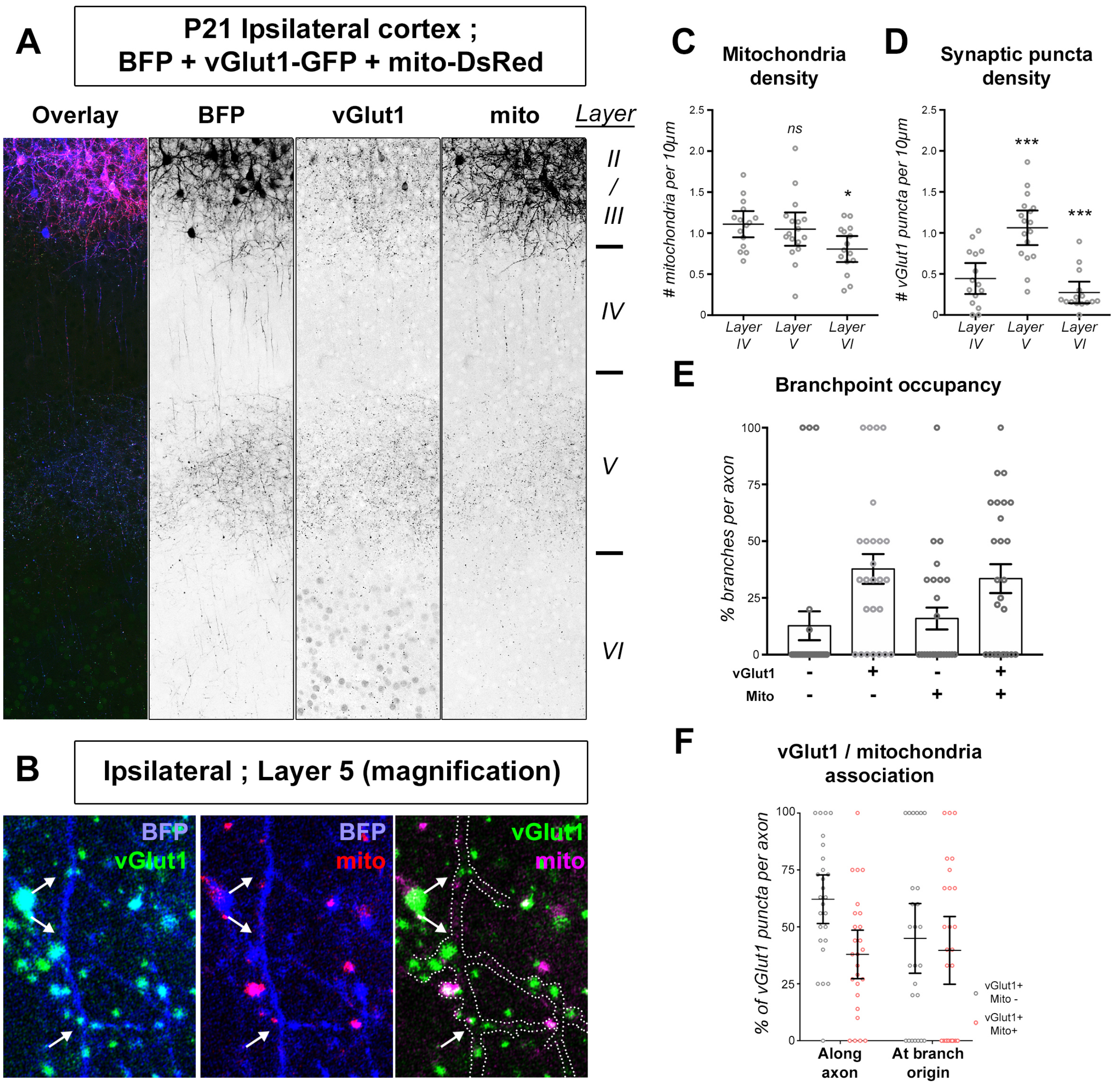


**Supplementary Figure 2 (related to Figure 2): distribution of axonal mitochondria and presynaptic boutons in vivo**

(A-B) Detail of the ipsilateral cortex of a mouse electroporated with plasmids coding the cell filler BFP, the presynaptic site marker vGlut1-GFP, and mito-DsRed. Magnification in layer V (B) shows individual axons and collaterals. White arrows point to branchpoints.

(C-D) Quantification of mitochondria and synaptic puncta density along individual axons. Each point represents the value for a given axon. Bars: median ± 95% CI. Statistical tests: one-way ANOVA with Tukey's multiple comparison (all data compared to Layer IV). $N_{(Layer\ IV)}=15$, $N_{(Layer\ V)}=17$, $N_{(Layer\ VI)}=15$.

(E) Quantification of branchpoint occupancy by either vGlut1+ puncta, mitochondria, or vGlut1+ puncta and mitochondria. Each point represents the value (%) for a given axon. Bars: average ± SEM. N=26 axons.

(F) Proportion of vGlut1 puncta devoid (grey) or associated to (red) mitochondria along the axon, and at branchpoints. Bars: median ± 95% CI. N=26 axons.

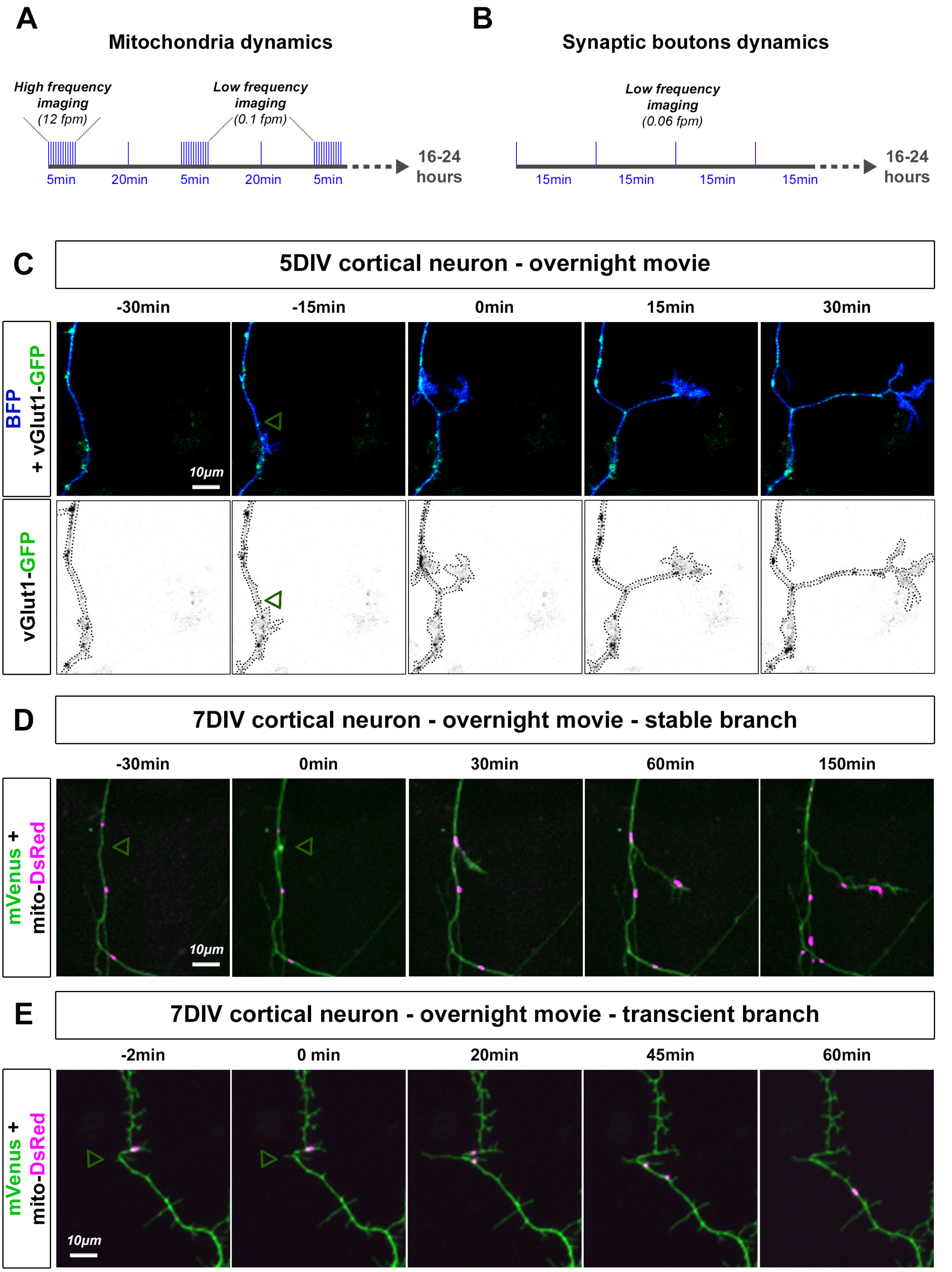


**Supplementary Figure 3 (related to Figure 3): dynamic correlation between presynaptic vesicle and mitochondria positioning and branch formation**

(A-B) Long-time imaging of axonal mitochondria and synaptic vesicles. 5-7DIV cortical neurons were imaged for 16-24 hours in cHBSS medium to visualize spontaneous events of branch formation. (A) For mitochondria, we alternated between pulses of high-frequency imaging (1 frame per 5 second, for 5 minutes) and low frequency imaging pauses (1 frame every 10 minutes for 10 minutes). (B) For synaptic vesicles, we imaged 1 frame every 15 minutes for 24 hours. Fpm: frame per minute.

(C) Distribution of vGlut1-GFP (green) before and after spontaneous branch formation in the axon of a 5DIV cortical PN. Future branch position is indicated by a green arrowhead. Note that future branch position is occupied by a vGlut1 puncta. BFP was used as a space filler.

(D-E) Examples of axons at 7DIV showing the distribution of mitochondria (magenta) before and after branch formation. Future branch position is indicated by a green arrowhead. Note the absence of mitochondria at the site of branch emergence prior to branch formation. mVenus was used as a space filler. (D) Stable branch typically recruited mitochondria at their origin and rapidly had mitochondria entering the branch. Eventually in some instances we could observe the pool of mitochondria at the branch origin moving into the branch. (E) On the contrary transient (unstable) branches failed to recruit and stabilize mitochondria at their origin

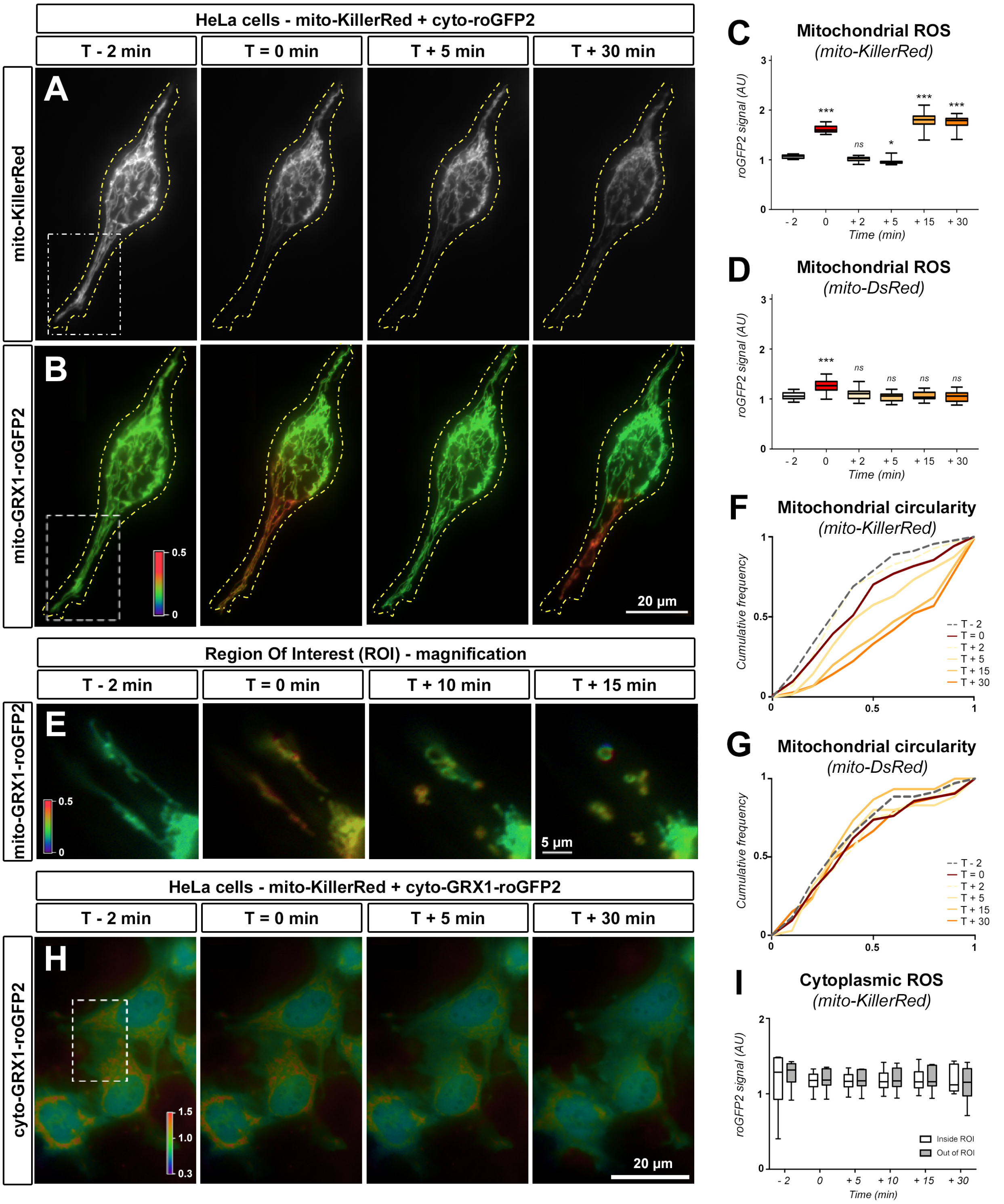


**Supplementary Figure 4 (related to figure 4): spatial and temporal validation of mitochondria photoinactivation using KillerRed**

(A-B) Example of a HeLa cell expressing mtKR (A) and mito-GRX1-roGFP2 (B) before and after photoinactivation of mitochondria in the ROI (white box). Laser intensity 100%, 30 seconds. Cell boundary indicated in yellow. mito-GRX1-roGFP2 is displayed as a fluorescence ratio between 405 and 488 illumination using a Rainbow LUT (Red: high ratio, oxidized. Blue: low ratio, reduced).

(C-D) Quantification of mitochondrial ROS accumulation (roGFP2 fluorescence ratio 405:488, normalized to T:-3 min, higher means more oxidized) at given time-points before, during and after CALI. Laser intensity 100%,

30 seconds. HeLa cells were co-transfected with mito-GRX1-roGFP2 and either mtKR or the control mito-DsRed, as indicated. Box-plot: 75$^{th}$ percentile, median and 25$^{th}$ percentile. Analysis: Repeated measures ANOVA with Dunnett's multiple comparison test. $N_{mtKR}$=11 cells, $N_{mtDsRed}$=29 cells.

(E) Magnification of the ROI of a HeLa cell before and after CALI showing mitochondria fragmentation and rounding.

(F-G) Mitochondria circularity was measured in the ROI before and after CALI in HeLa cells. Graphs show the cumulative circularity of mitochondria in cells expressing mtKR (F) or mtDsRed (G). N=9 cells.

(H) Representative image of HeLa cells co-transfected with mtKR and cyto-GRX1-roGFP2. roGFP2 signal is displayed as a fluorescence ratio between 405 and 488 illumination using a Rainbow LUT (Red: high ratio, oxidized. Blue: low ratio, reduced). CALI was performed in the indicated ROI (white box). Laser intensity 100%, 30 seconds.

(I) Quantification of cytoplasmic ROS accumulation by measuring cyto-GRX1-roGFP2 signal inside and out of the ROI showed no significant leakage of ROS in the cytoplasm of cells following photostimulation of mitochondria-targeted KillerRed. Box-plot: 75$^{th}$ percentile, median and 25$^{th}$ percentile. N=7 cells.

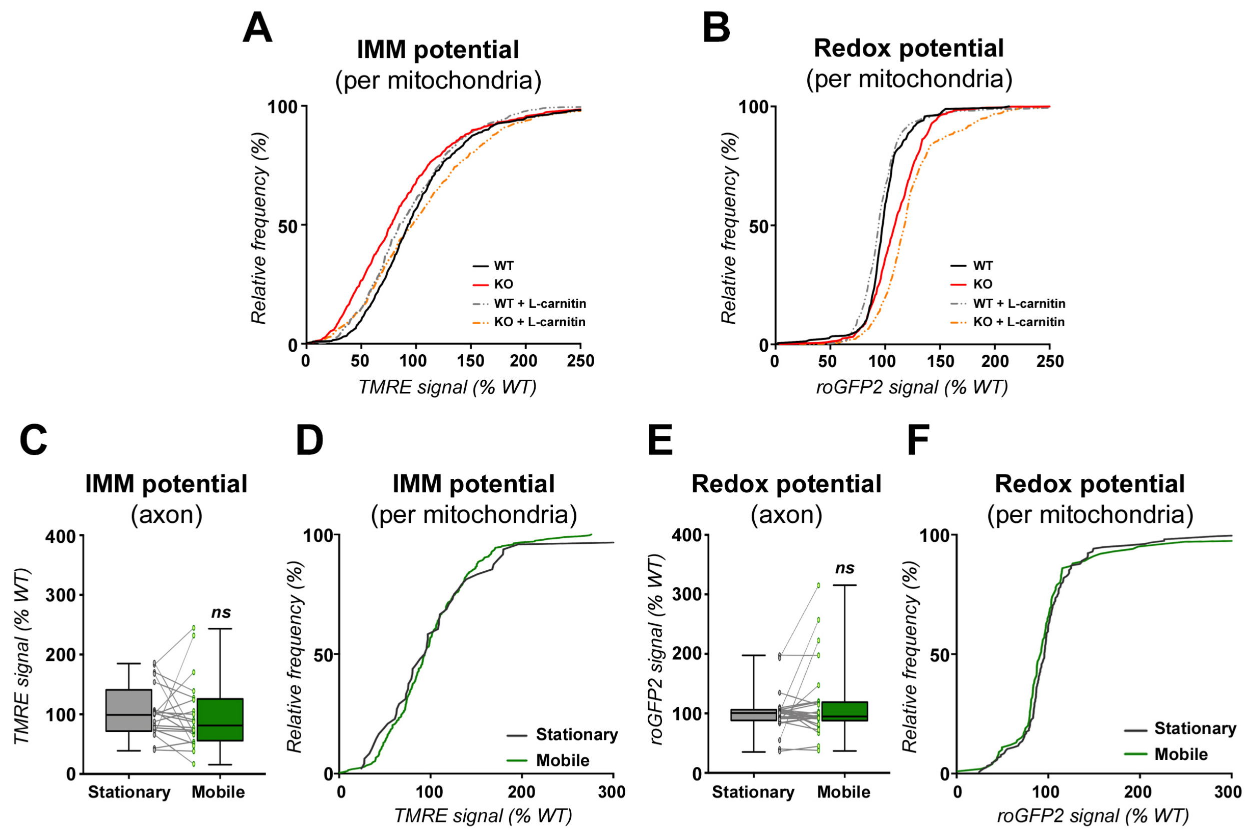


**Supplementary Figure 5 (related to figure 5): mitochondrial membrane potential and redox potential are not correlated with mitochondrial motility**

(A-B) Frequency distribution of individual mitochondria in the axon of NUAK1$^{F/F}$ neurons in conditions from Figure 5D-E. Plot of mitochondrial membrane potential (ΔΨm) (A) or redox potential (GRX1-roGFP2) (B).

(C-F) Effect of mitochondrial motility on IMM and redox potential in Wild-Type neurons. Time-lapse imaging was performed to sort out mobile (>5µm deplacement) or stationary mitochondria. (C, E) Graphs represent the average values for axonal mitochondria in a given axon. Box-plot: 75$^{th}$ percentile, median and 25$^{th}$ percentile. Individual dots represent average (per axon). Statistical tests: Wilcoxon matched-pairs ranked test. $N_{(TMRE)}$=20, $N_{(roGFP2)}$=26 individual neurons of 5 independent neuronal cultures. (D, F) Frequency distribution of individual mitochondria in the axon.

(G-J) Measurement of mitochondrial IMM and redox potential upon inhibition of the mitochondrial anchor SNPH to increase axonal mitochondrial motility. (G, I) Graphs represent the average values for axonal mitochondria in a given axon. Individual dots represent average (per axon). Statistical tests: Mann-Whitney. $N_{(TMRE,\ control)}$=15, $N_{(TMRE,\ control)}$=10 individual neurons of 5 independent neuronal cultures. $N_{(roGFP2,\ control)}$=27, $N_{(roGFP2,\ control)}$=24 individual neurons of 4 independent neuronal cultures. (H, J) Frequency distribution of individual mitochondria in the axon.

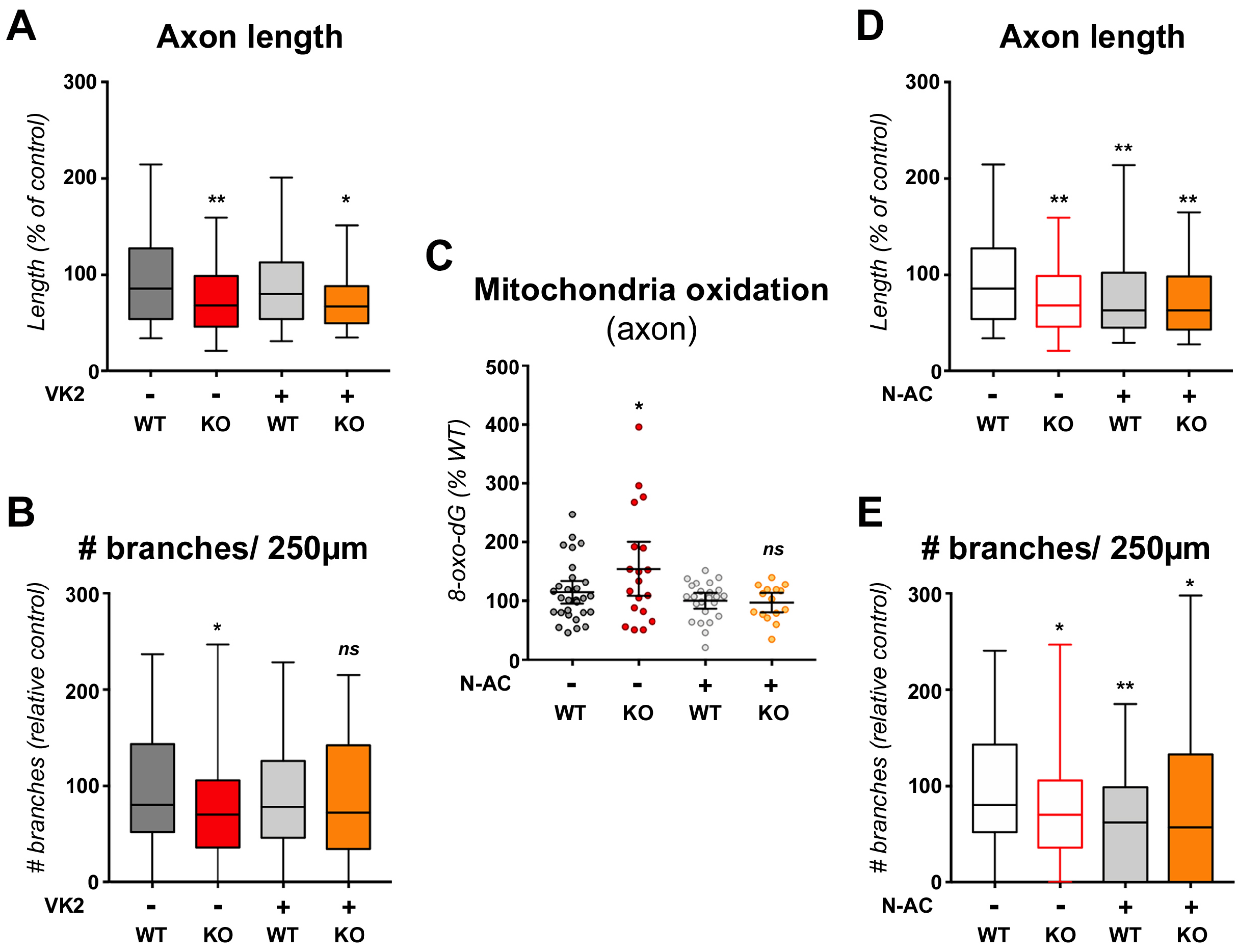


**Supplementary Figure 6 (related to figure 6): mitochondrial membrane potential and redox potential are not correlated with mitochondrial motility**

(A-B) Quantification of axon length and collateral branches of 5DIV neurons upon treatment with Vitamin K2 (VK2) (1mM) from DIV2 to DIV5. Box-plot: 75$^{th}$ percentile, median and 25$^{th}$ percentile. Statistical tests: Kruskal-Wallis test with Dunn's post-test (each condition compared to untreated WT condition). $N_{(WT)}$=202, $N_{(KO)}$=83, $N_{(WT+VK2)}$=163, $N_{(KO+VK2)}$=77 out of 3 independent experiments.

(C) Measurement of mitochondrial DNA oxidation by immunostaining with 8-hydroxy-2-deoxyGuanosine (8-oxo-dG) antibody. Fixed neuronal cultures at 5DIV were stained with 8-oxo-dG and neuronal fluorescence was measured as a readout of oxidative stress. For each field imaged, electroporated neurons (GFP positive) signal was normalized to non-electroporated neurons to account for variations in signal to noise. $N_{(WT)}$=29, $N_{(KO)}$=19, $N_{(WT+N\text{-}AC)}$=24, $N_{(KO+N\text{-}AC)}$=15 out of 12 independent fields (3 independent cultures).

(D-E) Quantification of axon length and collateral branches of 5DIV neurons treated with N-AcetylCysteine (N-AC) (1mM) from DIV2 to DIV5. Box-plot: 75$^{th}$ percentile, median and 25$^{th}$ percentile. Statistical tests: Kruskal-Wallis test with Dunn's post-test (each condition compared to untreated WT condition). WT and KO conditions are the same as (A-B). $N_{(WT)}$=202, $N_{(KO)}$=83, $N_{(WT+N\text{-}AC)}$=168, $N_{(KO+N\text{-}AC)}$=68 out of 3 independent experiments.
