## Supplementary material for "The AMPK-related kinase NUAK1 controls cortical axons branching though a local modulation of mitochondrial metabolic functions": Resources table

| REAGENT | SOURCE | IDENTIFIER (catalog number) |
|---|---|---|
| **Antibodies** | | |
| Mouse anti-Actin clone C4 | MP Biomedicals | 08691001 |
| Rabbit Anti-Citrate synthase (D7V8B) | Cell Signaling | 14309S |
| Rabbit anti-CRE recombinase (D7L7L) | Cell Signaling | 15036S |
| Chicken Anti-GFP Antibody | Rockland | 600-901-B12 |
| Hoechst 33258 solution | Merk/Sigma Aldrich | 94403 |
| Mouse Total OXPHOS Antibody Cocktail | Abcam | ab110413 |
| Rabbit Anti-TOM20 (D8T4N) | Cell signaling | 2406S |
| Donkey Anti-mouse HRP | Invitrogen | A16011 |
| Donkey Anti-Rabbit HRP | Invitrogen | A16029 |
| Goat Anti-Chicken AlexaFluor Plus 488 | Invitrogen | A32931 |
| Donkey Anti-Mouse AlexaFluor 647 | Invitrogen | A31571 |
| **Chemicals and drugs** | | |
| L-Carnitine (ISULIK 250ML) | Sogeval laboratories | 028AA308U |
| Rotenone | Merk/Sigma Aldrich | R8875 |
| D-Galactose | Merk/Sigma Aldrich | G0750 |
| Sodium Pyruvate solution | Merk/Sigma Aldrich | S8636 |
| N-Acetyl-L-cystein | Merk/Sigma Aldrich | A9165 |
| Menaquinone K4 | Merk/Sigma Aldrich | 47774 |
| Fast Green FCF | Merk/Sigma Aldrich | F7252 |
| Paraformaldehyde 32% | Electron Microscopy Sciences | 15714-S |
| Tetramethylrhodamine ethyl ester perchlorate (TMRE) | Merck, Sigma | 87917 |
| Potassium phosphate monobasic | Merk/Sigma Aldrich | P5655 |
| Disodium Succinate | Merk/Sigma Aldrich | 14160 |
| 2,6-Dichloroindophenol sodium salt hydrate (DCIP) | Merk/Sigma Aldrich | D1878 |
| Acetyl Coenzyme A sodium salt | Merk/Sigma Aldrich | A2056 |
| Oxaloacetic acid | Merk/Sigma Aldrich | O4126 |
| 5,5′-Dithiobis(2-nitrobenzoic acid) (DTNB) | Merk/Sigma Aldrich | D8130 |
| **Media and reagents for cell culture** | | |
| HEPES buffer; 1M; pH7,3 | Gibco | BP299 |
| D-glucose | Gibco | 47829 |
| HBSS (10X) | Gibco | 14185-045 |
| Papain, PDS kit, Papain Vial | Worthington | LK003176 |
| Deoxyribonuclease I from bovine pancreas | Merk/Sigma Aldrich | D5025 |
| Poly-D-lysine hydrobromide | Merk/Sigma Aldrich | P0899 |
| Laminin | Merk/Sigma Aldrich | L2020 |
| Neurobasal medium (1X) | Gibco | 21103-049 |

| | | |
|---|---|---|
| B27 serum free-supplement (50X) | Gibco | 17504-044 |
| N2 serum-free supplement (100X) | Gibco | A13707.01 |
| Glutamax supplement (100X) | Gibco | 35050-038 |
| Penicillin-Streptomycin solution (100X) | Gibco | 15140-122 |
| Neurobasal-A medium without D-glucose and sodium pyruvate | Gibco | A24775-01 |
| **Recombinant DNA** | | |
| pCAG-IRES-GFP (pCIG2) | Hand, R. et al. 2005 | NA |
| pCAG-CRE-IRES-GFP (pCIG2-CRE) | Hand, R. et al. 2005 | NA |
| pCIG2-shSNPH | Courchet et al. 2013 | NA |
| pCAG-mVENUS (pSCV2) | Hand and Polleux, 2011 | NA |
| pCAG-mitoDsRED | Courchet et al. 2013 | NA |
| pFUGW- PercevalHR | Tantama et al., 2013 | NA |
| pCAG-mito-KillerRed | This study | NA |
| pCAG-mScarlet-i | This study | NA |
| pCAG-cyto-Grx1-roGFP2 | Gift from Tommy L Lewis | NA |
| pCAG-mito-Grx1-roGFP2 | Gift from Tommy L Lewis | NA |
| pCAG-VGLUT1-Venus | Gift from Etienne Herzog | NA |
| pCAG-CRE | Gift from Seok-Kyu Kwon | NA |
| pCAG-mTagBFP2 | Lewis et al, 2016 | NA |
| pCAG-mito-mTagBFP2 | Lewis et al, 2016 | NA |
| pLKO.1 control | Courchet et al. 2013 | NA |
| pCAG | Courchet et al. 2013 | NA |
| **Other** | | |
| Glass bottom dishes | MatTek Corporation | P35G-1.5 |
| Borosilicate Glass capillaries O.D.:1mm,I.D.:0.50m, 10cm length, FiMT | World Precision Instrument | GBF100-50-10 |
